# Understanding the heterogeneity of alloreactive natural killer cell function in kidney transplantation

**DOI:** 10.1101/2023.09.01.555962

**Authors:** Dan Fu Ruan, Miguel Fribourg, Yuko Yuki, Yeon-Hwa Park, Maureen Martin, Geoffrey Kelly, Brian Lee, Ronaldo Miguel de Real, Rachel Lee, Daniel Geanon, Seunghee Kim-Schulze, Melissa McCarthy, Nicholas Chun, Paolo Cravedi, Mary Carrington, Peter S. Heeger, Amir Horowitz

## Abstract

Human Natural Killer (NK) cells are heterogeneous lymphocytes regulated by variegated arrays of germline-encoded activating and inhibitory receptors. They acquire the ability to detect polymorphic self-antigen via NKG2A/HLA-E or KIR/HLA-I ligand interactions through an education process. Correlations among HLA/KIR genes, kidney transplantation pathology and outcomes suggest that NK cells participate in allograft injury, but mechanisms linking NK HLA/KIR education to antibody-independent pathological functions remain unclear. We used CyTOF to characterize pre- and post-transplant peripheral blood NK cell phenotypes/functions before and after stimulation with allogeneic donor cells. Unsupervised clustering identified unique NK cell subpopulations present in varying proportions across patients, each of which responded heterogeneously to donor cells based on donor ligand expression patterns. Analyses of pre-transplant blood showed that educated, NKG2A/KIR-expressing NK cells responded greater than non-educated subsets to donor stimulators, and this heightened alloreactivity persisted > 6 months post-transplant despite immunosuppression. In distinct test and validation sets of patients participating in two clinical trials, pre-transplant donor-induced release of NK cell Ksp37, a cytotoxicity mediator, correlated with 2-year and 5-year eGFR. The findings explain previously reported associations between NK cell genotypes and transplant outcomes and suggest that pre-transplant NK cell analysis could function as a risk-assessment biomarker for transplant outcomes.

## INTRODUCTION

Human natural killer (NK) cells are a population of cytotoxic lymphocytes that play key roles in cancer surveillance, viral infection control, promoting normal pregnancy, and driving graft-versus-host disease and graft-versus-leukemia following hematopoietic stem cell transplantation. They are activated upon ligation of activating receptors concurrent with an absence of signaling between inhibitory receptors and class I HLA. NK cells undergo a developmental program of education instructed by interactions between NK cell receptors and class I HLA ligands. These interactions train NK cells to distinguish diseased cells with perturbed class I HLA expression and foreign cells with disparate class I HLA, from healthy cells with normal self class I HLA expression. Educated NK cells have greater functional capacity (cytokine production and cytotoxicity) to respond to reduced or, potentially, non-self HLA compared to immature and uneducated NK cells. In humans, interactions between class I HLA and NK cell receptors are determined by three genomic complexes, class I HLA which encode the A3, A11, Bw4, C1, and C2 ligands (chromosome 6), the natural killer complex (chromosome 12) which encodes the lectin-like CD94 and NKG2A, and the leukocyte receptor complex which encodes the killer-cell immunoglobulin-like receptors (KIR) (chromosome 19) ^1–3^. NKG2A interacts with HLA-E and educates a greater number of NK cells compared to KIR receptors ^4,5^. HLA-E cell surface expression is partly modulated by a dimorphism in the HLA-B leader peptide in which the -21 methionine (- 21M) dimorphism promotes greater stability in binding and folding by HLA-E and higher cell-surface expression compared to -21 threonine (-21T) leader peptides. Emerging evidence suggests that while the -21M dimorphism promotes higher HLA-E expression, its affinity to NKG2A may be lower compared to -21M leader peptides derived from HLA-A, -C, and -G ^6^. All HLA-C alleles and a minority of HLA-A and HLA-B alleles present epitopes that act as ligands for killer cell immunoglobulin-like receptors (KIR) which are encoded by highly polymorphic genes and vary in binding affinity to their cognate ligands ^3^. Due to the linkage disequilibrium of class I HLA alleles, HLA-B - 21M/T dimorphism, and the presence/absence of KIR ligands, HLA-A/B/C combine as haplotypes to discriminately educate diverse subsets NK cells across human populations ^7^.

Most studies acknowledge the underlying immunogenetics that regulate NK cell functions. However, NK cells are conventionally delineated as the CD56^bright^ cytokine-producing and CD56^dim^ cytolytic subsets, whereas transcriptome and mass cytometric (CyTOF) analyses have revealed that the combinatorial expression of many activating and inhibitory receptors in the NK cell repertoire yields a high diversity of subpopulations that have variable responses to stimulation ^8,9^. In the context of kidney transplantation, current concepts are that NK cell mediated antibody dependent cytotoxicity (ADCC) contributes to the effector mechanisms underlying antibody**-** mediated rejection (AMR). Deconvolution of microarray transcriptomic data has indeed found enrichment of NK cell transcripts in lesions defined pathologically as AMR ^10^. In one study, at least 25% of differentially expressed transcripts in kidney allografts in patients with detectable serum, donor-specific anti-HLA antibodies (DSA), and AMR associated with NK cells ^11^. These transcripts encoded proteins including granulysin, NKp80, and Ksp37, which suggest NK cell maturation and cytotoxic capacity. Most of these NK-associated transcripts are also enriched in patients who lack serum DSA but have histological evidence of AMR in allograft biopsies ^12^.

Whether and how NK cells function as antibody-independent, pathological mediators of kidney transplant injury is less well understood. Analyses of class I HLA and KIR genes in cohort studies of kidney transplant recipients suggest a role for NK cell education in mediating kidney transplant injury. A higher degree of mismatch between recipient KIR and donor KIR ligand-encoding HLA, termed “missing-self”, associates with a greater risk of microvascular inflammation ^13,14^. The incidence of chronic rejection and graft loss is also higher when there is missing-self or the absence of KIR2DL1/HLA-C2 and KIR3DL1/Bw4 interaction between donor and recipient ^15,16^. Beyond the observation that CD56^dim^ NK cells are elevated within allograft tissue from subjects with T-cell-medicated rejection (TCMR) ^17^, there is limited knowledge on the functional activity across specialized subsets of NK cells in kidney transplant recipients. Nor is there a clear understanding of how the immunogenetics that educate NK cells impact NK cell contributions to graft outcome.

To provide insight into antibody-independent pathologic functions of NK cells following kidney transplantation, we developed a panel of 44 antibodies to characterize peripheral blood-derived NK cell phenotypes and functions, pre- and post-transplantation, and before and after stimulation with donor cells, in kidney transplant recipients with defined outcomes from the Clinical Trials in Organ Transplantation-01 (CTOT01) and the CTOT19 cohorts ^18,19^.

## RESULTS

### NK cells vary in phenotype and composition across healthy donors and kidney transplant recipients

We performed CyTOF on PBMC of healthy individuals (n=20) and a subset of kidney transplant recipients from the CTOT01 cohort based on pre-transplant sample availability (n=70, see Methods) to characterize the heterogeneity of NK cells across subjects. CTOT01 was a previously described observational cohort of 280 kidney transplant recipients ^18^. As reported, the cohort included the following characteristics: 68.6% recipients of living donor kidneys, 28.6% African American, peak panel of reactive antibodies (PRA) was variable but less than 60% in 85.7% of the cohort, mean number of HLA (A, B, DR) mismatches was 3.5 out of 6, 44.6% received T cell depleting induction, and most subjects were maintained on tacrolimus, mycophenolate, and prednisone [see Table 1 in ^18^].

Unsupervised clustering ^20^ on the NK cell population yielded 18 subpopulations that we annotated into 9 primary subsets (Fig. 1, a-c, and Supplementary Figure 1). Consistent with our previous report ^9^, we identified both the CD56^dim^ and CD56^bright^ subsets with the CD56^dim^ subset comprising a mean of 87% of the NK cells. NKG2A^+^ NK cells were both CD56^bright^ and CD56^dim^ while KIR^+^ NK cells were primarily CD56^dim^. NK cell markers including CD56, CD57, and NKG2C displayed similar expression profiles in the healthy and CTOT01 kidney transplant cohorts. Most NK cell subpopulations observed in the healthy cohort were identified in the CTOT01 cohort in comparable proportions. Three NKG2A^+^ clusters, within the CD56^dim^NKG2A^+^KIR^+^ and CD56^bright^NKG2A^+^ subsets co-expressing activation markers IFNγ and XCL1, were higher in the CTOT01 cohort compared to healthy controls with a difference between 0.53% and 4.4% in the mean of cluster size across subjects (Fig. 1d). The CD56^−^CD16^+^CXCR3^high^CD27^high^ cluster was lower in the CTOT01 cohort (mean=0.51%) compared to the healthy cohort (mean=1.40%). All other subsets defined by unsupervised clustering were found in similar proportions between healthy donors and CTOT01 recipients.

**Fig. 1.**
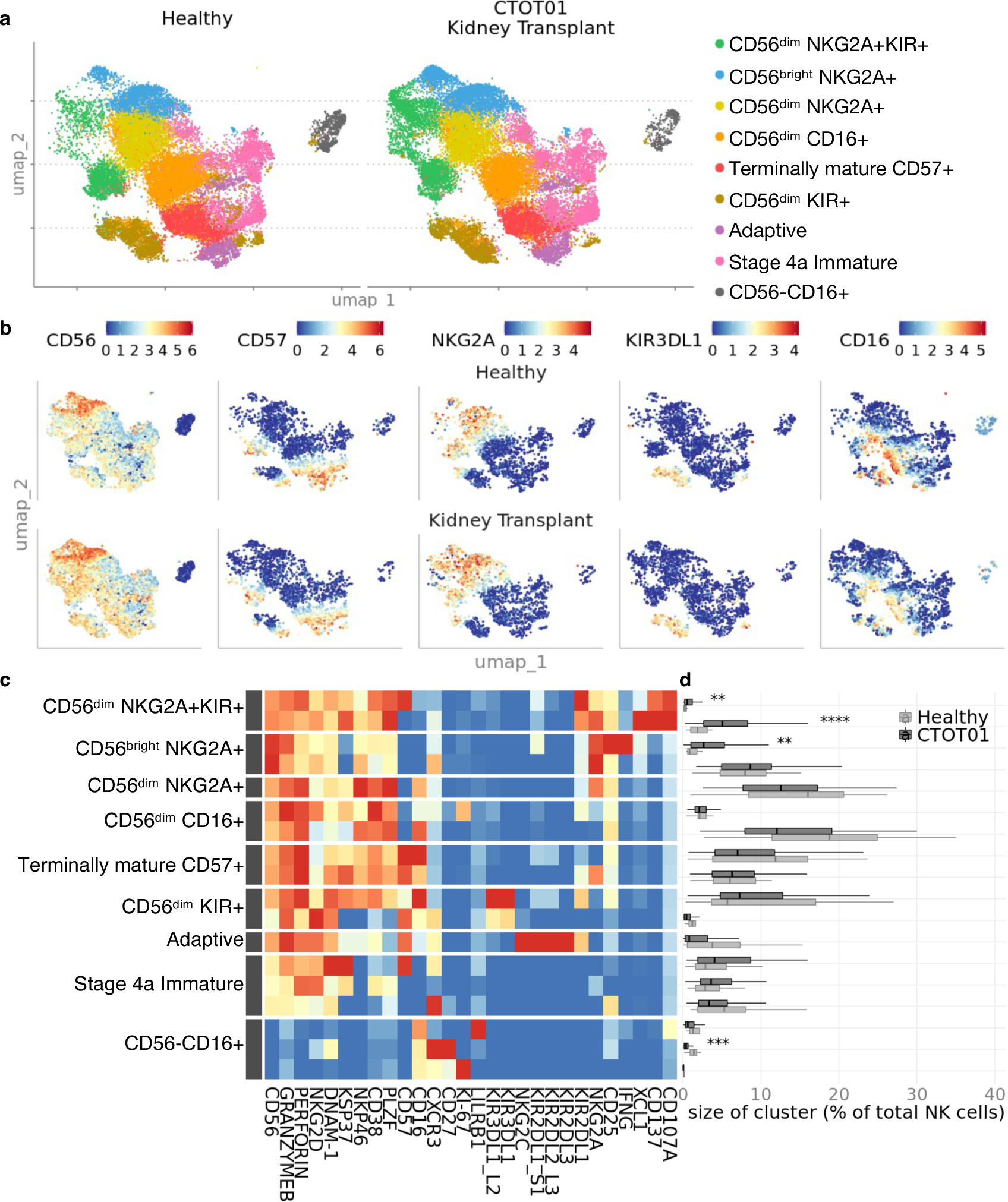
NK cells vary in phenotype and composition across healthy donors and kidney transplant recipients. Cyropreserved PBMC from healthy donor (n=20) and CTOT01 kidney transplant recipients (n=70) were recovered overnight in RPMI medium supplemented with 10 ng/mL recombinant human (rh) IL-15 and 10% heat-inactivated FBS. Results were profiled by CyTOF. **a** U-MAP of unsupervised RPhenograph clustering of peripheral blood derived NK cells shows heterogenous populations defined by variable expression of **b** NK cell markers CD56, CD57, CD16 and activating/inhibitory receptors such as NKG2A and KIR3DL1. **c** Heatmap shows median expression of CyTOF antibodies defining NK cell clusters. **d** Relative proportions of NK cell clusters across healthy donors and CTOT01 recipients were calculated by dividing number of cells in cluster by total NK cells per subject. *P* value was calculated using 2-sided unpaired Student’s *t*-test and values were adjusted for multiple testing with Bonferroni correction; \*\**p* < 0.01, \*\*\**p* < 0.001, \*\*\*\**p* < 0.0001.

To characterize the reactivity of NK cell subsets in response to allogeneic cells, we stimulated the healthy PBMC with wildtype K562 and HLA-E^+^ K562 cells, or with allogeneic non-transformed, peripheral blood-derived B cell lines (Fig. 2a) ^18^. We chose four allo-stimulator B cell lines to represent the presence/absence of the C1, C2, A03, A11, and Bw4 ligands in combination to maximize the number of KIR educating interactions to analyze. The allogeneic (allo) B cell stimulators were profiled for expression of activating NK receptor ligands by flow cytometry (Fig. 2b).

**Fig. 2.**
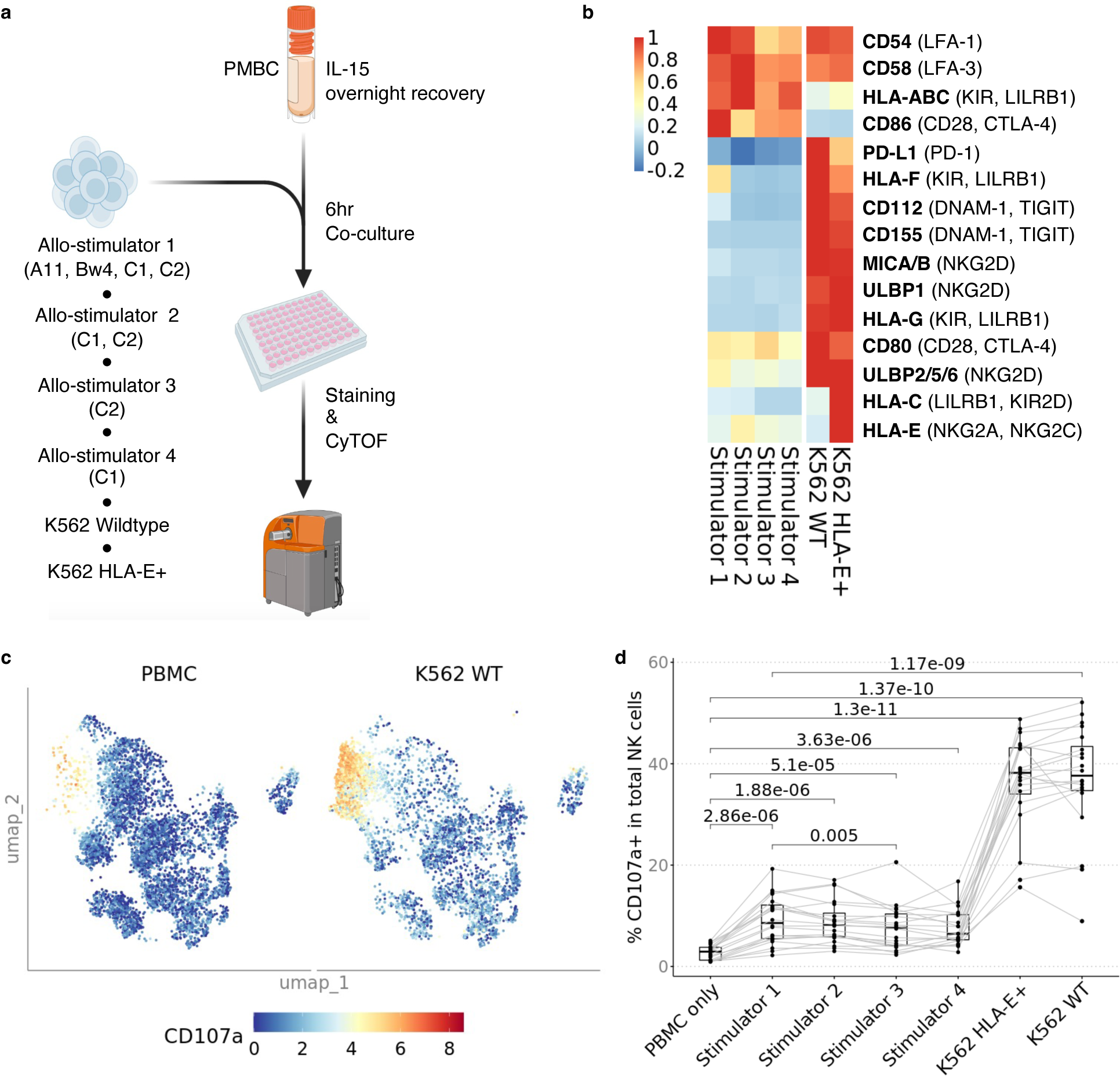
*In vitro* model of NK cell alloreactivity shows variability is dependent on donor ligand and recipient receptor differences. **a** Cryopreserved PBMC were recovered overnight in RPMI medium supplemented with 10 ng/mL rhIL-15 and 10% heat-inactivated FBS, and then stimulated with K562 wildtype (WT), K562 HLA-E+ and four allogeneic B cell lines expanded with CD40L-expressing fibroblasts and IL-4 at 3:1 E/T. **b** Heatmap showing MFI of NK cell ligands on stimulator cells; NK cell cognate receptor for ligands are indicated in parentheses. **c** U-MAP of CD107a expression in the coculture conditions PBMC only and PBMC co-cultured with wildtype K562. **d** Percentage of CD107a+ NK cells across healthy donors (n=20) when stimulated four allogeneic B cell stimulators, K562 WT and K562 HLA-E+. There were no differences between Stimulators 1-4 except between Stimulator 1 and Stimulator 3. All Stimulators induced less activation than K562 WT and K562 HLA-E+, and there was no difference between K562 WT and K562 HLA-E+. *P* value was calculated using 2-sided paired Student’s *t*-test and values were adjusted for multiple testing with Bonferroni correction.

Data from studies of NK cell reactivity in virus infection and cancers indicate that non-educating activating ligands including adhesion molecules CD54 and CD58 bind LFA-1 and LFA-3 respectively. MICA/B and ULBP1/2/5/6 bind the activating NKG2D receptor ^21^. Other expressed ligands, including CD112 and CD155, were shown to be stimulatory if bound to activating receptor DNAM-1, but suppressive when bound to inhibitory receptor TIGIT ^22^. Similarly, class I HLA molecules may be activating or inhibitory depending on the NK cell receptor bound ^23^. Based on these published findings from other systems, we predicted that the alloreactive response of a given NK cell population depends on the combination of ligands on stimulator cell and the combination of receptors on the NK cell. The four allo-stimulator B cell lines we used expressed a similar profile of NK cell activating ligands while the two K562 cell lines had increased expression of additional NK ligands including ULBP1, ULBP2/5/6 and MICA/MICB. When we stimulated NK cells with the allo-stimulator cells or with the K562 cell lines, they increased expression of degranulation and activation markers CD107a, XCL1, IFNγ, and CD137. We observed the strongest responses to the K562 cell lines which lack HLA-ABC expression and highly express activating ligands CD155 and MICA/MICB (Fig. 2c, d, and Supplementary Fig. 2a-c). To deepen our understanding of how various NK cell subsets contribute to this overall response, we focused on specific subsets based on maturation and education.

### Mapping the rules that govern NK cell allo-reactivity

Immature human NK cells are characterized by CD56^bright^ expression and are potent producers of cytokines and chemokines ^24^. They lack expression of perforin and granzymes A/B, restricting their role to production of cytokines and chemokines that shape their microenvironment. CD56 expression is downregulated as NK cells mature; these mature CD56^dim^ NK cells have greater cytotoxic potential than CD56^bright^ NK cells. During maturation, NK cells continue to increase cytotoxic potential and decrease cytokine production. These NK cells can be identified by expression of CD16, KIRs, and the terminal maturation marker CD57. Separately, educated NK cells are CD56^dim^ and express NKG2A and/or self-class I HLA-reactive KIR and are capable of detecting aberrations in class I HLA expression ^7^. To study the effect of education, we gated on CD56^dim^ NK cells and further defined subpopulations based on CD57 and the educating receptors NKG2A and KIR2DL1/S1, KIR2DL2, KIR2DL3, KIR3DL1, and KIR3DL2. KIR^+^ NK cells, which are educated through the KIR/HLA-A/B/C axis, were defined as expressing any of KIR2DL1/S1, KIR2DL2, KIR2DL3, KIR3DL1, or KIR3DL2 independent of KIR ligands encoded on class I HLA. For both CD57^+^ and CD57^−^ NK cells, expression of the educating receptors associated with greater percent positivity of CD107a in response to K562 cells; we observed the same effect in education in response to the four allo-stimulator cell lines expressing combinations of the A11, Bw4, C1, and C2 KIR ligands (Fig. 3a, Supplementary Fig. 3).

**Fig. 3.**
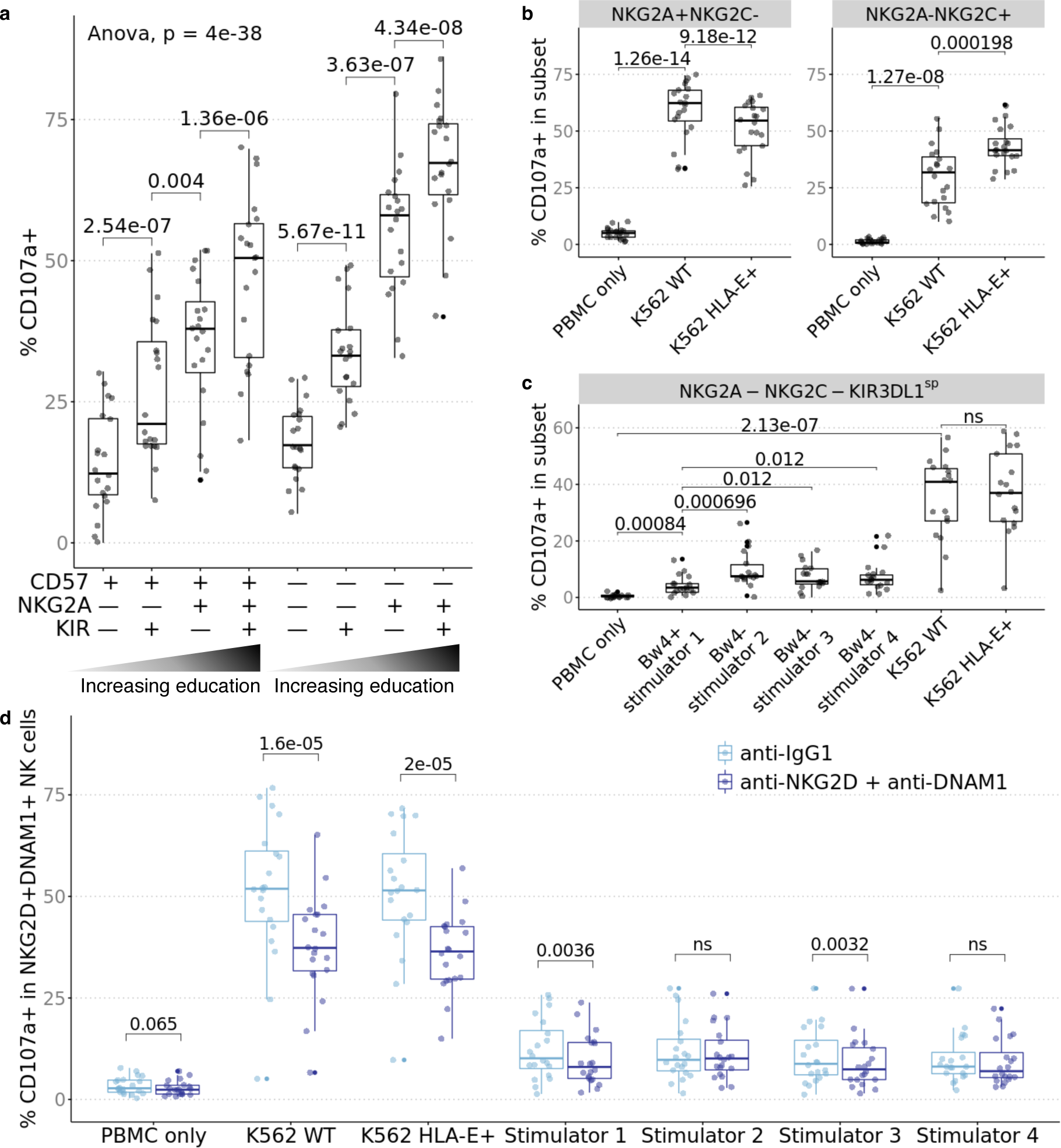
Educated NK cell subsets are more responsive to missing-self and NK cell activation can be abrogated by blockade of activating receptors. PBMC from healthy donors (n=20) were cocultured with stimulators and profiled by CyTOF. **a** Subsets were defined by gating on CD56^dim^ NK cells followed by combinatorial expression of educating inhibitory receptors, NKG2A, KIR3DL1, KIR3DL2, KIR2DL1/S1, KIR2DL2 and KIR2DL3 and the CD57 maturation marker. Percent CD107a^+^ when stimulated by K562 WT increases with NKG2A/KIR expression. **b** NKG2A^+^NKG2C^−^ NK cells increased CD107a when stimulated by K562 WT and increase was depressed by stimulation with HLA-E+ K562. NKG2A^−^NKG2C^+^ NK cells increased CD107a when stimulated by K562 which was further increased by stimulation with HLA-E+ K562. **c** In individuals who expressed *KIR3DL1* and class I HLA alleles that encode Bw4 (n=17), NKG2A^−^NKG2C^−^KIR3DL1^sp^ NK cells increased CD107a in response to 4 allogeneic stimulator B cells and K562 but was relatively inhibited by stimulator 1 which uniquely expressed Bw4. **d** Percent CD107a^+^ for NKG2D^+^DNAM-1^+^ NK cells was decreased by blockade of NKG2D and DNAM-1. *P* value was calculated as one-way ANOVA as indicated and as 2-sided paired Student’s *t*-test; *t*-test *p* values were adjusted for multiple testing with Bonferroni correction and ns indicates *p* > 0.1.

NK cell subsets expressing inhibitory NKG2A, but lacking the activating isoform, NKG2C, i.e., NKG2A^+^NKG2C^−^ NK cells, were activated by wildtype K562 cells. And as predicted, the same NK cell subset responded less to HLA-E^+^ (inhibitory ligand) K562 cells. Conversely, we observed augmented degranulation of CD107a by NKG2A^−^ NKG2C^+^ stimulated with HLA-E^+^ K562 cells (Fig. 3b). To characterize the effect of Bw4 education, we gated on KIR3DL1^+^NKG2A^−^NKG2C^−^ NK cells in individuals who express *KIR3DL1* and class I HLA alleles that encode Bw4 (n=17). This subset degranulated in response to all stimulators but there was no difference between stimulation by wildtype or HLA-E^+^ K562, neither of which express the cognate ligand Bw4 ligand (Fig. 3c). Amongst the 4 allo-stimulator cell lines, allo-stimulator 1, which uniquely expressed Bw4, specifically and dominantly induced weaker responses by KIR3DL1^+^NKG2A^−^ NKG2C^−^ NK cells, consistent with it functioning as an inhibitory ligand.

To test whether other activating interactions (that do not influence the education program ^23^) affect NK cell reactivity against allogeneic cells, we repeated these cocultures in the presence/absence of blocking antibodies to these activating receptors. NK cell activation induced by missing-self and non-self was abrogated by blockade of non-educating activating receptors such as NKG2D and DNAM-1 (Fig. 3d). The reduction in CD107a degranulation by NKG2D^+^DNAM-1^+^ blockade was greatest when the NK cells were co-cultured with wildtype and HLA-E^+^ K562 which highly expressed ligands for NKG2D and DNAM-1 (e.g. CD112, CD155, MICA/B, and ULBP). Together we conclude that interactions via NKG2D and DNAM-1 contribute to the alloreactive NK cell responses and that blockade of these interactions specifically abrogates activation in the subsets that express the activating receptors.

### Circulating NK cell phenotypes are stable over time following transplantation

We characterized the diversity within the repertoire of peripheral blood-derived NK cells in the CTOT01 kidney transplant cohort prior to transplantation and at 6-to 24-months post-transplantation, in subjects with available post-transplant PBMC (n=36). CyTOF analysis and unsupervised clustering revealed that the relative proportions of NK cell subsets remarkably did not differ pre-transplant (on dialysis) vs. post-transplant, despite induction and maintenance immunosuppression (Fig. 4a-c).

**Fig. 4.**
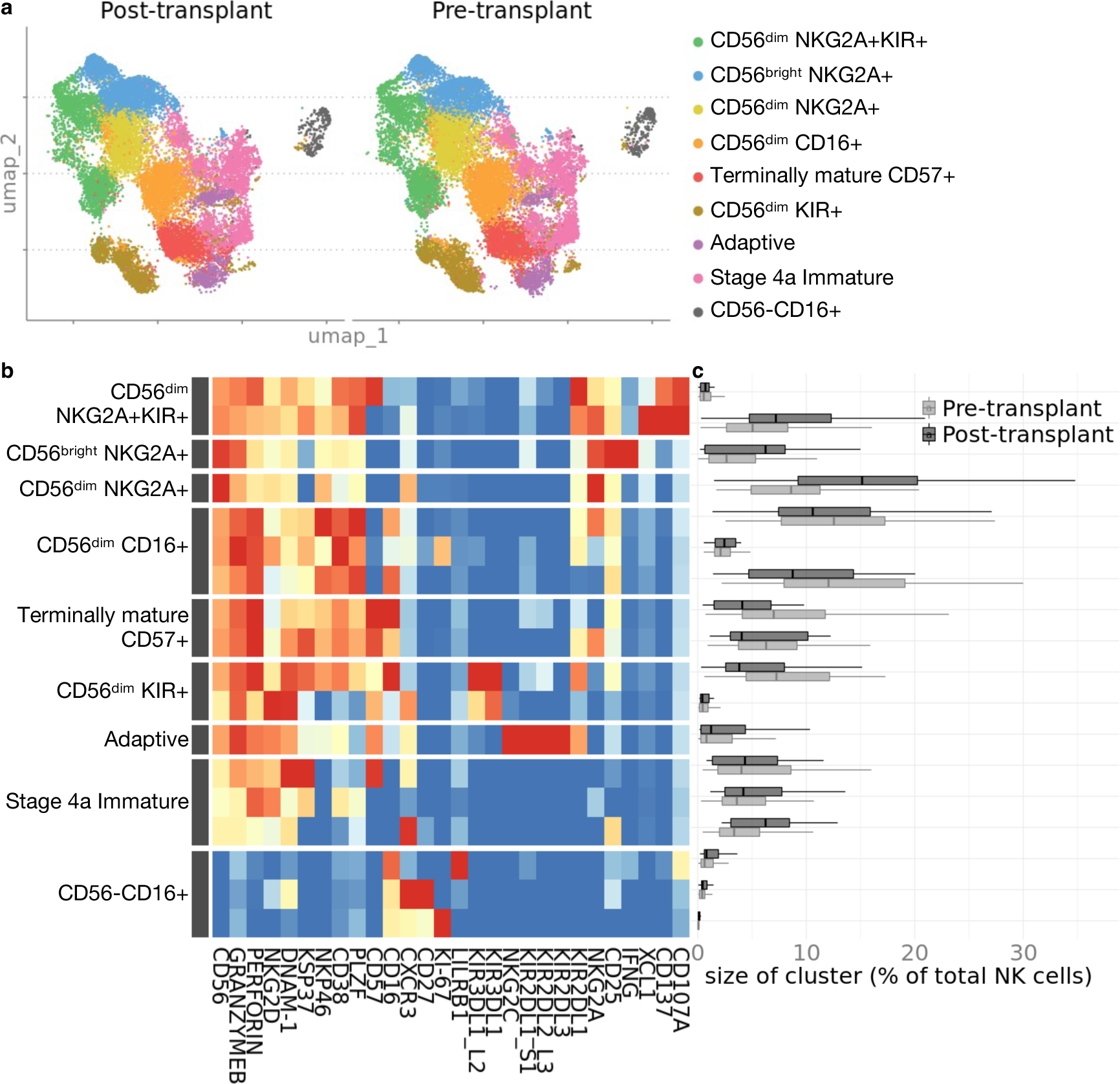
Phenotype and composition of NK cells are stable pre- and post-transplant. Cyropreserved PBMC from CTOT01 kidney transplant (n=70) were recovered overnight in 10 ng/mL IL-15 and profiled by CyTOF. **a** U-MAP projection of unsupervised RPhenograph clustering of CTOT01 recipient pre-transplant (n=70) and post-transplant (n=36) peripheral blood derived NK cells. **b** Heatmap shows median expression of CyTOF markers defining NK cell clusters. **c** Proportions of NK cell subsets in pre-transplant and post-transplant *ex vivo* samples were calculated by dividing number of cells in cluster by total NK cells per sample. There were no differences in proportional size of clusters between pre- and post-transplant. *P* value was calculated using 2-sided unpaired Student’s *t*-test and values were adjusted for multiple testing with Bonferroni correction.

To determine whether the observed effects of education on NK cell reactivity in the healthy cohort were conserved in transplant recipients, we stimulated CTOT01 PBMC obtained pre-transplant and 6-18 months post-transplant with their allogeneic donor B cells. Analysis of the pre-transplant samples showed, despite heterogeneity of NK ligands expressed on donor allo-stimulator cells, significantly increased CD107a degranulation in NK subsets expressing either or both NKG2A and KIR, suggesting that the effect of education transcends donor differences (Fig. 5). As in the healthy cohort, we gated on CD56^dim^ NK cells and KIR^+^ NK cells were defined as expressing KIR2DL1/S1, KIR2DL2, KIR2DL3, KIR3DL1, or KIR3DL2 independent of the recipient’s educating KIR ligands. In an additional analysis, we also refined the KIR^+^ populations for each recipient to exclude any KIR^+^ NK cell that did not have cognate KIR ligand encoded on the recipient’s class I HLA (Supplementary Fig. 4). For all subsets analyzed, including educated NKG2A^+^ and KIR^+^ subsets, CD107a expression was equivalent between pre- and post-transplant. Educated subsets released more CD107a than uneducated subsets pre- and post-transplant. We also observed a similar pattern amongst subsets for pre- and post-transplant expression of XCL1, CD137 and IFNγ, including higher expression by NKG2A and/or KIR-expressing NK cells (Supplementary Fig. 5). While levels of CD107a, XCL1 and CD137 were maintained post-transplant, the percentage of IFNγ producers was lower for the CD57^+^NKG2A^−^KIR^−^ and CD57^+^NKG2A^−^KIR^+^ subsets post-transplant. However, the decrease in percentage of IFNγ producers was only 1.4% and 1.9% for the CD57^+^NKG2A^−^KIR^−^ and CD57^+^NKG2A^−^KIR^+^ subsets respectively compared to the 12.1% to 13% of IFNγ producers in the CD57^−^ NKG2A^+^KIR^−^ CD57^−^NKG2A^+^KIR^+^ NK cell subsets. Overall, we found that NK cell subsets expressing either or both NKG2A and KIR were more responsive to transplant donor cells and that this alloreactivity persisted post-transplant.

**Fig. 5.**
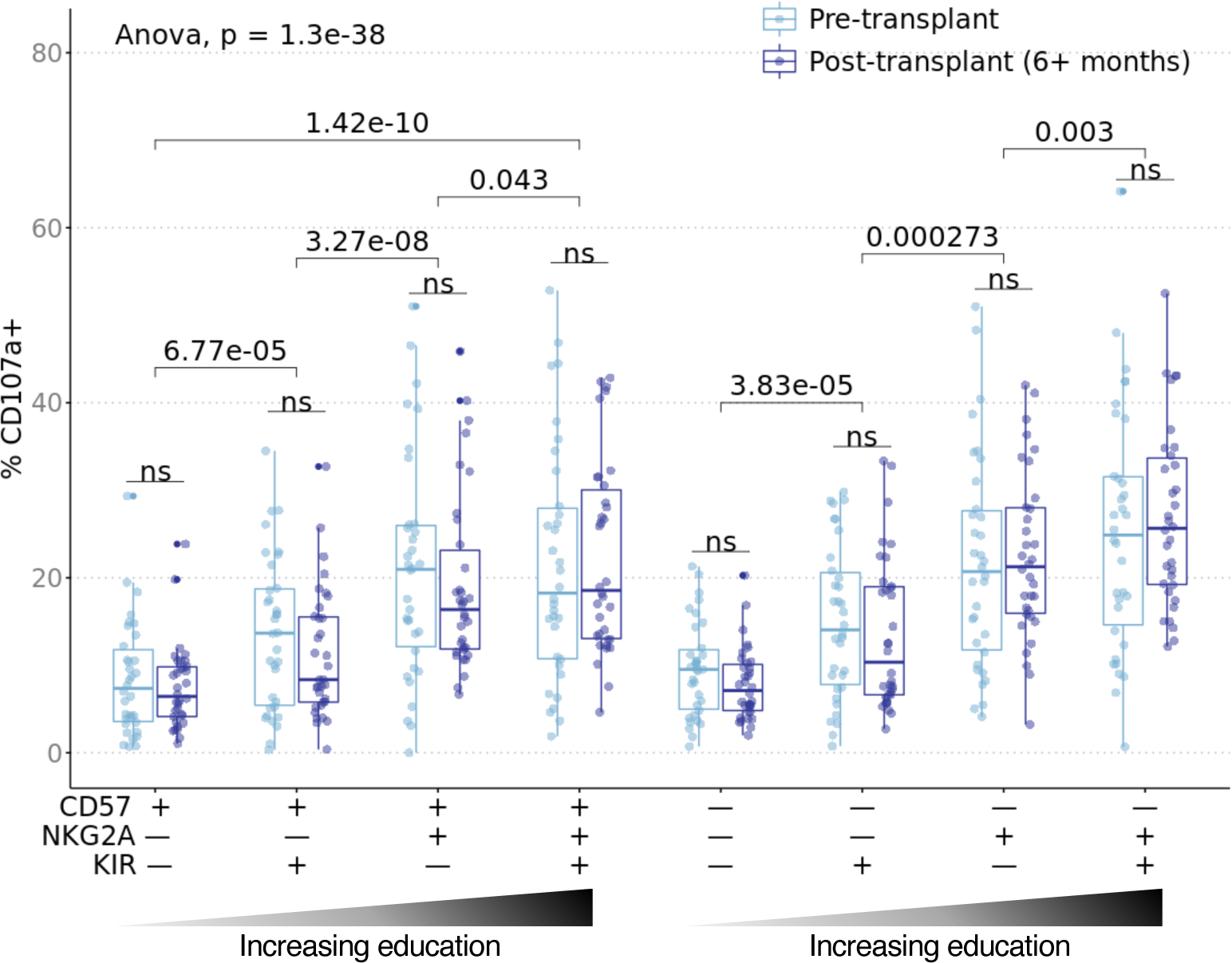
Alloreactivity of educated NK cells transcend donor differences and is maintained post-transplant. Cryopreserved pre- and post-transplant PBMC from CTOT01 kidney transplant (n=70) were recovered overnight in 10 ng/mL rhIL-15 and stimulated with donor allo-stimulator B cells for 6 hours at 3:1 E/T. Results were profiled by CyTOF and subsets were defined by gating on CD56^dim^ NK cells followed by combinatorial expression of educating inhibitory receptors, NKG2A, KIR3DL1, KIR3DL2, KIR2DL1 and KIR2DL3 and the CD57 maturation marker. Boxplot shows percent CD107a^+^ in recipient NK cell subsets in response to co-culture with donor cells for which pre- and post-transplant cells were available (n=34). When recipient NK cells were stimulated with donor cells, the NKG2A^+^KIR^+^ educated subset produced the most CD107a while the uneducated NKG2A^−^KIR^−^ subset produced the least CD107a. The effect of NKG2A/KIR education on CD107a expression in response to donor cells persisted 6+ months post-transplant. *P* value was calculated as one-way ANOVA as indicated and as 2-sided paired Student’s *t*-test; *t*-test *p* values were adjusted for multiple testing with Bonferroni correction and ns indicates *p* > 0.1.

As a further test of the functional relevance of NKG2A/HLA-E interaction in inhibiting allogeneic NK cells, we quantified HLA-E surface expression on donor cells by flow cytometry. We stratified donor cells in the top 75th^th^ percentile as having high HLA-E and the remaining as low HLA-E. We analyzed the effects of high and low HLA-E expression on NKG2A^+^NKG2C^−^ NK cell function. Recipient NKG2A^+^NKG2C^−^ NK cells stimulated by donor cells with higher HLA-E (n=16) expression produced more XCL1 and degranulated/released more granzyme B than recipient NKG2A^+^NKG2C^−^ NK cells stimulated by donor cells with lower HLA-E (n=51) (Fig. 6a-c, Supplementary Fig. 6).

**Fig. 6.**
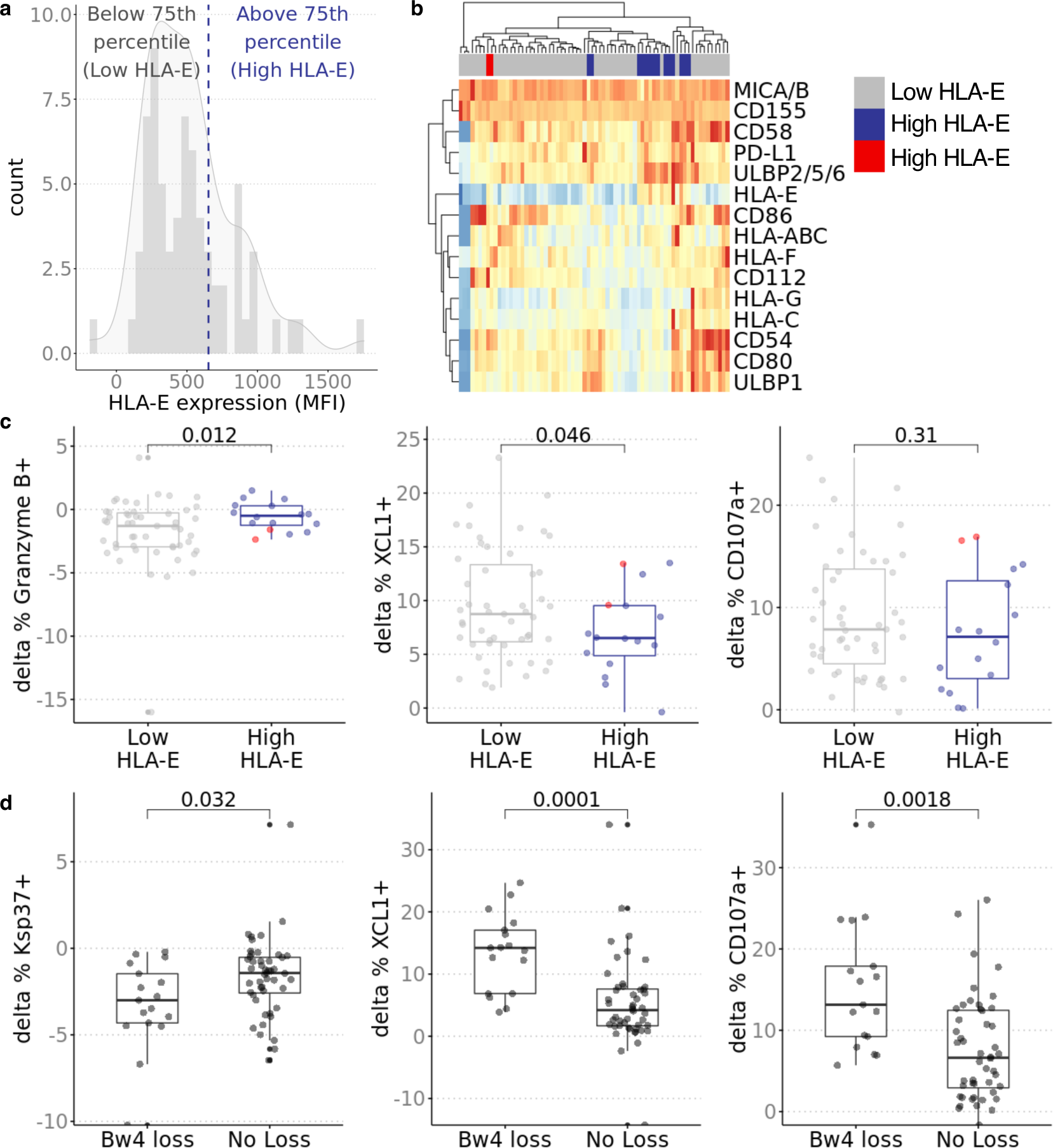
Effects of NKG2A and KIR3DL1 education apply in transplant donor/recipient pairs. **a** Distribution of HLA-E expression in donor stimulator cells. HLA-E expression on donor cells was measured in triplicate by flow cytometry. Median fluorescence intensity was normalized by isotype control. **b** Stimulator groups were defined by HLA-E expression where 75th percentile was high (n=51) and remaining were low (n=16). Hierarchical clustering shows similar expression profile across HLA-E low donors with exception of two indicated in red in heatmap legend. These two donors expressed higher CD112 and HLA-F; additional analysis excluding these two donors show stronger associations (Supplementary Fig. 5). **c** Change in percent of XCL1^+^ and Granzyme B^+^ (delta between stimulated and PBMC only condition) in recipient NKG2A^+^NKG2C^−^ NK cells was greater when stimulated by donors with lower HLA-E (n=16) than donors with higher HLA-E (n-51). Two donors with higher CD112/HLA-F (red dots) induced more Granzyme B, XCL1 and CD107a production in NKG2A^+^NKG2C^−^ NK cells compared to other donors with high HLA-E expression. **d** Bw4 loss was defined as lower Bw4 copy number in transplant donor compared to recipient. No loss of Bw4 was defined as equal or greater copy number of Bw4 on HLA-A/B alleles in donor compared to recipient or recipient was *KIR3DL1*^−/−^. When gating on KIR3DL1^+^ NK cells, percent of CD107a^+^ and XCL1^+^ and degranulation of Ksp37 was higher when there was loss (n=17) than when there was no loss (n=48). *P* value was calculated using 2-sided unpaired Student’s *t*-test; nominal *p* values shown.

To determine the effect of KIR3DL1/Bw4 education on alloreactivity, we categorized donor/recipient pairs based on copies of Bw4 encoded by their HLA-A and HLA-B alleles. Whereas missing-self reactivity has been traditionally defined as complete lack of the ligand, in the transplant setting, donor cells may have lower copy number of Bw4 rather than absence of Bw4. We categorized those with loss of Bw4 as recipients paired with donors with fewer copies of Bw4 (n=17). Recipients without loss of Bw4 had donors with equal or greater copy number of Bw4 or do not express KIR3DL1 (n=48). Within the pre-transplant KIR3DL1^+^ NK subset, we found that those with Bw4 loss produced more CD107a and XCL1 compared to those without loss of Bw4 (Fig. 6d). We also analyzed the release of Ksp37, a secretory protein expressed by cytotoxic lymphocytes, that has been associated with inflammatory states such as asthma and severe Covid-19 ^25,26^. In our analysis, percentage of Ksp37 staining in the KIR3DL1^+^ NK subset was decreased in recipients with Bw4 loss compared to recipients without Bw4 loss suggesting that the loss of Bw4 inhibition on KIR3DL1^+^ NK cells promoted the release of Ksp37. Collectively, our data suggest that education contributes to NK cell alloreactivity in kidney transplant recipients and that the effect of education is not significantly diminished by commonly used immunosuppression regimens that are highly effective at suppressing T cells.

### Allo-induced Ksp37 release correlates with NK cell cytotoxicity

Next, we investigated the killing potential of activated NK cells in our *in vitro* assays. We flow-sorted NK cells from three healthy blood donors for their uneducated NKG2A^−^KIR^−^ population and the NKG2A^−^KIR^+^, NKG2A^+^KIR^−^, and NKG2A^+^KIR^+^ populations. KIR^+^ cells were identified by positive staining with either anti-KIR3DL1/L2 or KIR2D. To avoiding the confounding effects of NKG2C/HLA-E interactions, we excluded all NKG2C^+^ cells were excluded from these comparisons. We stimulated these NK cells with wildtype K562 and allo-stimulator cells and analyzed stimulator cell death and NK cell CD107a and Ksp37 expression by flow cytometry (Supplementary Fig. 7). Wildtype K562 and allo-stimulator cell death increased when co-cultured with NK cells compared to stimulator cell alone (Fig. 7a, b). The analyses showed that NKG2A^+^ NK cells induced more cell death than NKG2A^−^KIR^−^ NK cells with some heterogeneity amongst the three healthy donors. Across these NK subsets, the degree of wildtype K562 and allo-stimulator cell death correlated with lower intracellular Ksp37 (implying release) and CD107a degranulation (Fig. 7c, d). The majority of Ksp37 loss, CD107a degranulation, and stimulator cell death was dominantly mediated by the educated NK cell subsets that express NKG2A and/or KIR. Together with the data on Ksp37 release in KIR3DL1^+^ NK cells from recipients with loss of donor Bw4 inhibition, our data suggest Ksp37 functions as a sensitive marker of NKG2A^+^ and KIR^+^ NK cell killing and activation.

**Fig. 7.**
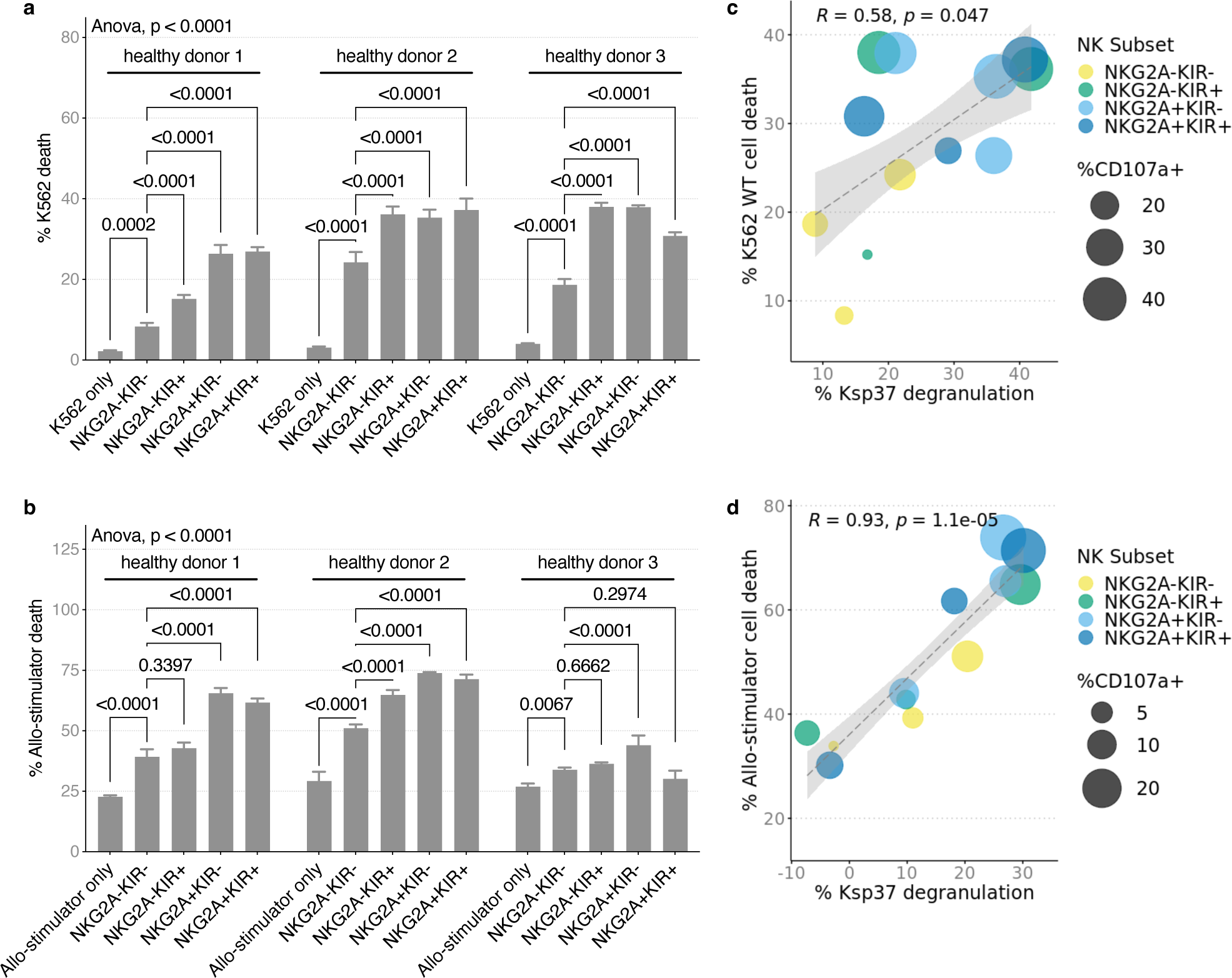
Educated NK cell subsets are more allo-reactive and are potent killers. Healthy donor PBMC were isolated from buffy coat and rested overnight in 10 ng/mL IL-15. NK cell subsets were flow sorted and co-cultured with K562 WT and all-stimulator B cells at 1:1 E/T for 5 hours to assess NK cell killing and activation. **a** K562 WT cell death and **b** allo-stimulator cell death were increased by addition of NK cells in culture. Error bars show standard deviation of mean on technical replicates. **c** Ksp37 release by NK cells was positively correlated with percent of K562 WT and **d** allo-stimulator cell death. High cell death and Ksp37 release was dominated by educated NK cell subsets and high CD107a expression. *P* value in **a** and **b** was calculated using two-way ANOVA as indicated, and as 2-sided unpaired Student’s *t*-test adjusted by Tukey’s multiple comparisons correction. *P* value in **c** and **d** are calculated by Pearson’s correlation.

### Pre-transplant NK cell reactivity to donor cells correlates with late allograft function

Since missing-self as determined by HLA/KIR genotyping independently associates with microvascular injury (MVI) and similar numbers of NK cells have been detected in MVI^+^DSA^+^C3d^−^ and MVI^+^DSA^−^ biopsies ^14,27^, we tested the hypothesis that pre-transplant donor-reactive NK cell Ksp37 degranulation correlates with post-transplantation kidney allograft function. We defined eight NK cell subsets based on combinatorial expression of CD57, NKG2A, and KIR and correlated pre-transplant Ksp37 release in response to donor cells within each subset with recipient estimated glomerular filtration rate (eGFR) at several timepoints post-transplantation. In the CD57^−^ NKG2A^+^KIR^−^ subset, lower frequency of Ksp37-expressing cells correlated inversely with eGFR at two years and at five years post-transplantation, independent of induction immunosuppression, thymoglobulin induction, delayed graft function, rejection status, living/deceased status of transplant donor, and degree of HLA mismatch (Fig. 8a, Supplementary Table 1a-b, and Supplementary Table 2a-b). Effect of DSA on this association is unlikely (but cannot be formally ruled out) as there were only two subjects with detectable serum DSA. This observation was unique to Ksp37 in the CD57^−^ NKG2A^+^KIR^−^ subset as combinations of other subsets and other functional markers (IFNγ, CD107a, XCL1, and CD137) did not correlate with 2- or 5-year eGFR. To validate this observation, we repeated the allo-stimulation co-cultures using donor/recipient pairs from the deceased donor kidney transplant CTOT19 cohort ^19^. These assays showed a significant decrease in percentage of Ksp37-expressing CD57^−^NKG2A^+^KIR^−^ NK cells (consistent with release of Ksp37) that also correlated inversely with 6-month and 2-year eGFR (n=25) (Fig. 8b). The inverse correlation between Ksp37 release and eGFR was independent of delayed graft function in the CTOT19 validation cohort, but this cohort was underpowered to additionally correct for acute rejection (n=2) and the number of HLA-A/B/C of mismatches (Supplementary Table 3a-c and Supplementary Table 4a-c).

**Fig. 8.**
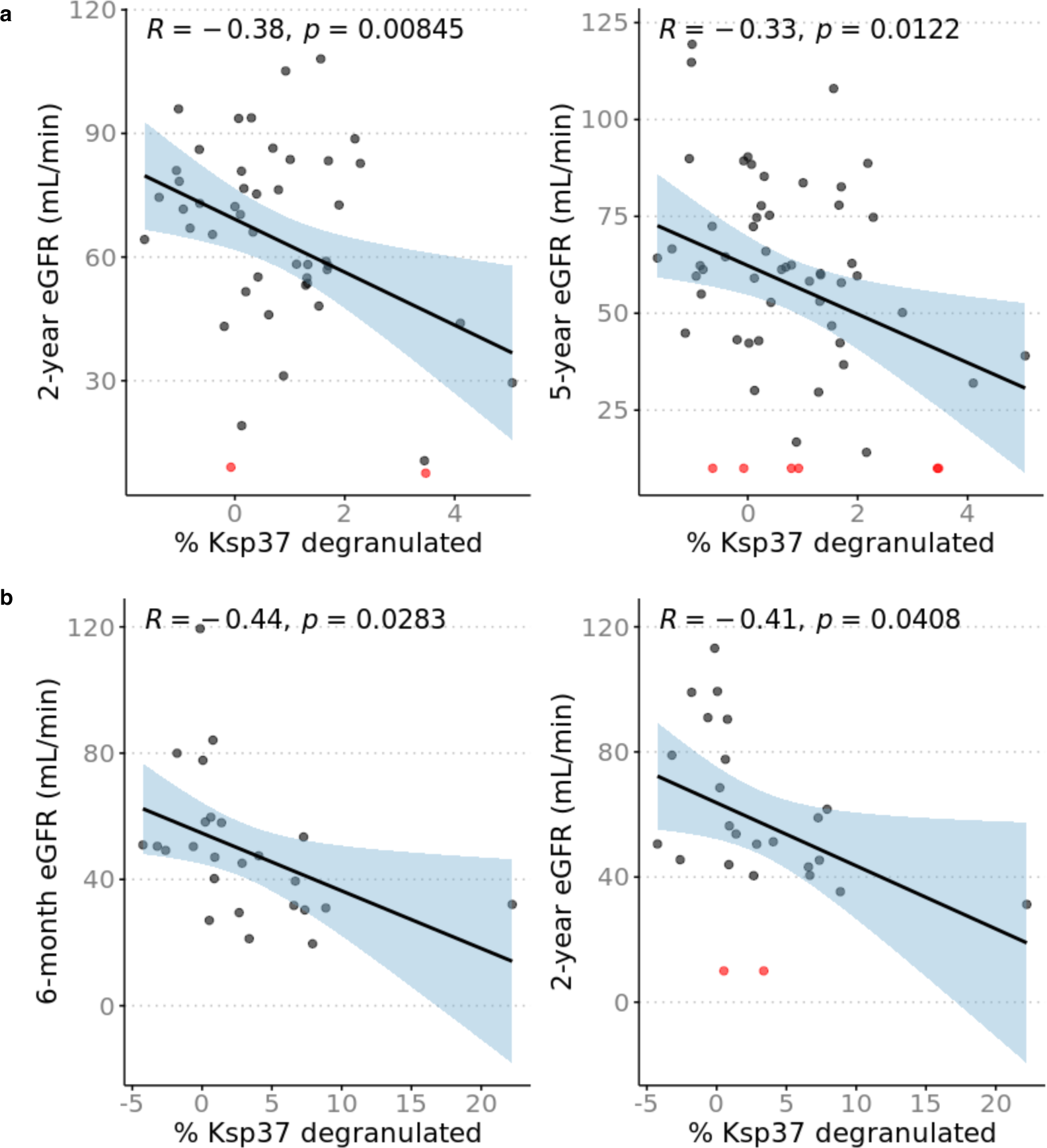
Increased Ksp37 degranulation in NK cells correlate with lower estimated glomerular filtration rate (eGFR). Percent Ksp37^+^ calculated as percent Ksp37^+^ NK cells in B cell coculture condition minus PBMC only condition. eGFR values were calculated with CKD-EPI formula and patients with graft failure were assigned eGFR = 10 mL/min. **a** Greater degranulation and decrease of %Ksp37^+^ in CD57^−^NKG2A^+^KIR^−^ NK cells correlates with higher 2-year eGFR (n=47) and 5-year eGFR (n=58) in CTOT01. **b** Greater degranulation and decrease of %Ksp37+ in CD57^−^NKG2A^+^KIR^−^ NK cells correlates with higher 6-month and 2-year eGFR (n=25) in validation cohort CTOT19. Patients with eGFR ≤ 10 mL/min are indicated in red. Correlation coefficient and *p* value shown for Pearson correlation.

## DISCUSSION

Although NK cells have been found to play critical roles in controlling cancers and microbial infections, the transplantation field lacks a nuanced understanding of how diverse NK cell subsets regulated by class I HLA, KIR, and NKG2A-mediated education and variegated co-expression of activating receptors contribute adversely or therapeutically. Previous studies have demonstrated that NK cells and KIR genetics relating to NK cell activity associate with kidney transplant outcomes independent of T cells and DSA. While the increase in CD16 seen in AMR has suggested ADCC as the mechanism by which NK cells mediate graft injury, emerging indirect genetic evidence suggests that there are alternate ways by which alloreactive NK cells contribute to long-term allograft outcomes beyond AMR ^28,29^. One such alternate way is missing-self reactivity, which necessitates an understanding of the HLA-I/KIR/NKG2A immunogenetics that inform NK cell education ^13,14,17,27^. In this study, we analyzed HLA/KIR/NKG2A immunogenetics in donors and recipients alongside high-dimensional profiling of recipient-derived NK cell subsets to elucidate the mechanisms by which variably educated NK cell subsets contribute to long-term graft function.

Mass cytometric analyses of peripheral blood-derived NK cells revealed a high degree of heterogeneity and showed that this diversity is consistent between healthy and pre-transplantation kidney transplant recipients. Interestingly, clustering analyses designated three *ex vivo* NK cell clusters with basal levels of activation (IFNγ^+^, CD137^+^, CD107a^+^, and/or XCL1^+^) that were elevated in the kidney transplant cohort pre-transplant. Two of these clusters were CD56^dim^NKG2A^+^KIR^+^ and one was CD56^bright^NKG2A^+^; whether these populations found in the periphery share a common lineage as the kidney-resident NK cells other groups have detected in transplant biopsies remains to be studied.

All peripheral and most tissue-derived CD56^bright^ NK cells express NKG2A, which might frame the HLA-E/NKG2A axis as an explanation for the prevalence of CD56^bright^ NK cells seen in rejection biopsies of previously published studies ^8,30^. Consistent with these findings, the CD57^−^NKG2A^+^KIR^−^ population in our transplant cohorts released Ksp37 that associated with poor long-term graft function. The donor-induced release of Ksp37 in our *in vitro* models also offers a mechanism to explain the detection of this transcript in prior studies that developed gene signatures of chronic injury in kidney transplantation ^11,12,31^. Since our model for predicting eGFR is independent of acute rejection, it bolsters emerging evidence that NK cells contribute to graft injury regardless of rejection status. In particular, it might explain why higher degree of missing-self as determined by recipient and donor genotype has been associated with MVI, which does not meet the BANFF criteria for rejection ^27^.

We profiled in a cohort of healthy individuals the rules of NK cell alloreactivity to non-self and how this process is regulated by class I HLA, KIR and NKG2A. *In vitro* models of PBMCs and allogeneic stimulator cell co-cultures identified significant correlation between the breadth of NK cell activation with their education status, i.e., NKG2A- or KIR3DL1-educated NK cells were uniquely inhibited by their cognate HLA-E and Bw4 ligands, respectively, and dominantly activated relative to non-educated subsets when in absence of their inhibitory ligands. We also showed that within healthy donors of different genetic backgrounds, NKG2A/KIR expression promotes alloreactivity against all stimulator cell lines. NK cells that co-expressed NKG2A and KIR were more alloreactive than NK cells that expressed either NKG2A or KIR. We demonstrated that the system of education is preserved and engaged in transplant patients and that the magnitude of alloreactivity transcends differences amongst transplant donor cells as seen by a similar pattern of alloreactive response in pre-transplant NK cells against donor cells. The increase in KIR3DL1^+^ NK cells expressing CD107a and XCL1 and degranulating Ksp37 in recipients stimulated by donors deficient in Bw4 or with lower copies of Bw4 provide a mechanism for the genetic association studies linking the absence of recipient KIR3DL1 and donor Bw4 to the incidence of chronic rejection ^15^.

We also showed that the effects of education on alloreactivity persist post-transplant after NK cells have been exposed to induction immunosuppression and continuing exposure to maintenance immunosuppression. The two most common types of induction immunosuppression, rabbit antithymocyte globulin (rATG) and IL-2 receptor antagonists (IL2RA), are aimed at depleting T cells (rATG) or reducing the T cell proliferation (IL2RA) and have demonstrated efficacy in reducing early acute rejection ^32^. Maintenance immunosuppression following kidney transplantation mostly consists of calcineurin inhibitors such as cyclosporin A and tacrolimus, mTOR inhibitors, and mycophenolate, which have been shown to reduce rejection by reducing T cell proliferation and activation. *In vitro* ADCC assays on peripheral blood-derived NK cells show that Cyclosporine A and tacrolimus reduces IFNγ and CD107a at high doses while mycophenolate and sirolimus have minimal effects on IFNγ and CD107a ^33^. However, their effect on NK alloreactivity in kidney transplantation is less well understood. Recent studies characterizing the effects of mTOR and calcineurin inhibitors on NK cell functions in solid organ transplantation have presented discordant results. There is some evidence suggesting that NK cells from kidney transplant recipients vary expression of activation markers depending on the type of immunosuppression administered and that post-transplant NK cells from calcineurin-treated recipients produce less cytokines compared to healthy individuals ^34,35^. Yet, other studies show in murine models of kidney transplantation that calcineurin inhibitors fail to prevent NK cell-mediated rejection while mTOR inhibitors may reduce NK cell missing-self-induced microvascular injury ^27,36^.

Our results do not address whether post-transplant NK cells are less alloreactive compared to healthy donor NK cells in response to the same allogeneic stimulator cells. Rather, we show that NKG2A/KIR-educated NK cells are more alloreactive than uneducated NKG2A^−^KIR^−^ NK cells and that this enhanced donor-induced alloreactivity is maintained post-transplant meaning that maintenance immunosuppression is insufficient to address this facet of alloreactivity. Although there is a small decrease in the abundance of post-transplant circulating IFNγ^+^ NK cells compared to pre-transplant, it is restricted to the less educated NKG2A^−^KIR^−^ and NKG2A^−^KIR^+^ subsets.

We show that NK cell alloreactivity can be mitigated *in vitro* through the blockade of their activating receptors NKG2D and DNAM-1. Studies in various settings of solid organ transplantation point to the relevance of NKG2D and its ligands in promoting graft injury. In murine models, NKG2D expression increases over the course of ischemic injury and that this injury is reduced by the adoptive transfer of *NKG2D*^−/−^ NK cells or through blockade of NKG2D ^37^. Similarly, NKG2D blockade has been shown to reduce cardiac vasculopathy in murine heart transplantation ^38,39^. Although there is less evidence for the role of NKG2D and DNAM-1 in solid organ transplantation, blockade of these two receptors in hematopoietic stem cell transplantation for AML decreased degranulation of NK cells against allogeneic T cells ^40^. Our CyTOF analyses of circulating NK cells show that NKG2D and DNAM-1 are co-expressed on educated Ksp37-degranulating subsets that may be contributing to injury of kidney allografts; thus, there is potential for targeting these two activating receptors, amongst other activating pathways, to reduce kidney injury.

Future studies to visualize NK cell subsets localized to areas of immune infiltrate or fibrosis may clarify the helper or direct roles of NK cells in mediating graft injury. The results of the current study provide an in-depth analysis of the heterogeneity of NK cells found in the periphery of transplant recipients and how education and expression inhibitory/activating receptors guide their alloreactive response. We provide an explanation for the published associations between genotype and outcomes and raise the possibility that pre-transplant NK cell analyses could be used as risk assessment biomarkers for kidney transplantation.

## METHODS

### Cohorts and subjects

CTOT01 was a prospective multicenter observational trial of kidney transplants from 2006-2009 led by Drs. Heeger and Hricik. Trial design and details on subjects have been previously published ^18^. CTOT19 was a randomized control trial to test Infliximab as induction therapy in deceased donor kidney transplantation from 2016-2021 led by Drs. Heeger and Hricik. Trial design and details on subjects have been previously published ^19^. The CTOT01 (n=70) and CTOT19 (n=26) subjects studied herein were chosen based on sample availability. Values for eGFR were extracted from the primary publications. CTOT01 2-year and 5-year eGFR values were calculated with the Chronic Kidney Disease Epidemiology Collaboration (CKD-EPI) formula for adults and the Schwartz formula for children ^18,41^. CTOT19 6-month and 24-month eGFR values used in this study were calculated with the CKD-EPI equation ^19^. Patients with graft failure were assigned eGFR = 10 mL/min. PBMC from the healthy cohort (n=20) were sourced as buffy coats from the New York Blood Center (New York, NY, USA) and from the Bhardwaj Lab (Mount Sinai, New York, NY).

### HLA and KIR Typing

DNA was extracted from frozen whole blood, buffy coats, cryopreserved PBMC, and cryopreserved B cells using the DNeasy Blood & Tissue Kit (Qiagen, cat # 69506). The Carrington Lab (National Cancer Institute, Bethesda, MD) and CD Genomics (Shirley, NY) performed class I HLA and KIR typing (Supplementary Tables 5-7).

### Profiling stimulator cell lines

Allo-stimulator B cells were obtained from previous study where transplant donor primary B cells were expanded using CD40L-transfected fibroblasts and IL-4 according to previously published protocol ^18,42^. Wildtype and HLA-E+ K562 cell lines were provided by Deepta Bhattacharya. All stimulator cells were profiled for activating and inhibitory ligands by staining with a viability dye and surface antibodies (Supplementary Table 8) in FACS buffer (DPBS, 2% heat-inactivated FBS, 2 mM EDTA) for 30 minutes on ice; cells were stained in triplicate with isotype controls. After washing, cells were fixed with 2% paraformaldehyde (PFA) (EMS, cat #15710) for 10 minutes at room temperature, resuspended in FACS buffer, and stored at 4°C until acquisition on LSRFortessa (BD Biosciences). Results were analyzed on Cytobank software (Beckman Coulter) and R software version 4.0.3.

### *In vitro* cocultures

Transplant recipient and healthy donor cryopreserved PBMC were thawed and recovered overnight in 10 ng/mL recombinant human IL-15 (Peprotech, cat #200-15) at 2*10^6^ cells/mL cell culture media (RPMI-1640, 10% heat-inactivated fetal bovine serum (FBS), 1% Penicillin, 1% Streptomycin, 1% L-glutamine). Allo-stimulator B cells were thawed immediately prior to coculture. After overnight recovery, PBMCs were cocultured with donor allo-stimulator B cells, K562 wildtype or K562 HLA-E+ cell lines at 3:1 E/T for 6 hours in cell culture media supplemented with anti-CD107a-172Yb (Miltenyi, cat #130-124-536) in 96-well round-bottom plates. Plates were centrifuged at 1000 rpm for 2 minutes at start of coculture. Following 1 hour of incubation, Brefeldin (Biolegend, cat #420601) and Monensin (BioLegend, cat #420701) were added for remainder of coculture. For NKG2D/DNAM-1 blockade in the healthy cohort, PBMCs were incubated with cocktail of anti-NKG2D-145Nd (BioLegend, cat #320814) and anti-DNAM-1-146Nd (Life Technologies, cat #MA5-28149) or mouse IgG1 κ isotype control (BioLegend, cat #401402) for 30 minutes at room temperature prior to addition of stimulator cells. Assays were ended by placing plate on ice for 10 minutes and proceeding to CyTOF staining protocol.

### CyTOF staining

Antibodies were either purchased in metal-conjugated form from Fluidigm or purchased as purified carrier-free antibodies and conjugated following Fluidigm’s Maxpar Labelling protocols. Antibodies were titrated, and master mixes of cocktails were prepared prior to experiments. Technical replicates from cocultures were pooled and washed in cell staining media (PBS, 0.2% BSA). Samples were Fc-blocked (BioLegend, cat #422302) on ice for 5 minutes and stained with anti-β2m Platinum (Pt) barcodes (Human Immune Monitoring Core, Mount Sinai, NY) for 30 minutes on ice. 194Pt, 195Pt, and 196Pt-barcoded samples were pooled for surface staining with antibody cocktail (Supplementary Table 9) and Rh103 (Fluidigm, cat #201103A) for 30 minutes in room temperature. Surface antibody cocktail for healthy cohort samples blocked with anti-NKG2D-145Nd and anti-DNAM-1-146Nd in culture excluded the staining antibodies for anti-NKG2D and anti-DNAM-1. After surface staining, CTOT01 and healthy donor cocultures were fixed and permeabilized following manufacturer protocol using eBioscience™ Foxp3 / Transcription Factor Staining Buffer Set (ThermoFisher, cat #00-5523-00). For CTOT19 coculture, cells were fixed and permeabilized following manufacturer protocol using Cytofix/Cytoperm™ Fixation/Permeabilization Kit (BD Biosciences, cat #554714). After centrifugation and washing, samples were stained with Cell-ID 20-Plex Pd Barcoding kit (Fluidigm, cat #201060) for 30 minutes in room temperature. Barcoded samples were then pooled for intracellular staining for 30 minutes on ice with antibody cocktail supplemented with Heparin 100 U/mL. After washing, cells were fixed and Ir-stained with 2.4% PFA, 1% saponin, and 0.05% Ir (Fluidigm, cat #201192A). Sample of all pooled cells were stored at -80°C in 10% DMSO/FBS until acquisition.

### CyTOF analyses

Live intact NK cells were identified by gating in CytoBank, and events were exported for analysis in R software version 4.0.3 (Supplementary Fig. 1). A maximum of 2000 events from each subject/experimental condition were randomly selected for all downstream unsupervised clustering analyses. All events for live intact NK cells were included for manual gating analyses. Pre-processing of FCS files and events for analysis was done with cytoqc version 0.99.2 and MetaCyto version 1.12.0. Expression values of each marker were Arcsinh transformed with cofactor of 5. Unsupervised clustering was performed with R package Rphenoannoy version 0.1.0, and dimensionality reduction for generating UMAPs was done with R package umap version 0.2.10.0. Additional manual gating and Boolean gating to define NK subpopulations were done using flowCore version 2.2.0 and premessa version 0.3.2. Hierarchical clustering and visualization of heatmaps were generated with pheatmap version 1.0.12.

### NK cell killing assay and flow cytometry analyses

PBMC from healthy donor buffy coats were isolated by Ficoll density centrifugation and rested overnight in 10 ng/mL recombinant human IL-15 (Peprotech, cat #200-15) at 2*10^6^ cells/mL cell culture media (RPMI-1640, 10% FBS, 1% Penicillin, 1% Streptomycin and 1% L-glutamine). After overnight recovery, PBMCs were enriched for NK cells by negative selection following manufacturer protocol (Stemcell Technologies, Cat #19055) and surfaced stained in FACS buffer (DPBS, 5% heat-inactivated FBS, 2 mM EDTA) with antibody cocktail of CD3, CD56, NKG2A, NKG2C, KIR3DL1/L2, and KIR2D (Supplementary Table 10) for 30 minutes on ice. Stained cells were washed and resuspended in sorting buffer (RPMI-1640, 2% heat-inactivated FBS, 1% Penicillin, 1% Streptomycin, 1% L-glutamine, 1:3000 propidium iodide) prior to sorting on Cytoflex SRT (Beckman Coulter Life Sciences). Stimulator cells were stained with CFSE 1:33000 following manufacturer protocol (Life Technologies, cat #C34570). Sorted NK subsets were cocultured with CFSE-stained stimulator cells at 1:1 E/T in cell culture media supplemented with anti-CD107a in 96-well V-bottom plates. Plates were centrifuged at 1000 rpm for 2 minutes at start of coculture. After 5 hours, assay is ended by placing plate on ice for 10 minutes before proceeding to viability staining with Zombie NIR and surface staining as detailed above. Surface-stained cells were fixed with 2% PFA for 10 minutes at room temperature, permeabilized (BioLegend, cat #421002) for 30 minutes on ice, and stained with intracellular anti-Ksp37 for 30 minutes on ice. Cells were fixed with 2% PFA for 10 minutes at room temperature, resuspended in FACS buffer, and stored at 4°C until acquisition on LSRFortessa (BD Biosciences). Results were analyzed using Cytobank and R software version 4.0.3.

### Illustrations and graphs

Graphical abstract and Figure 2a was created with BioRender.com. Graphs were generated with GraphPad Prism or R package ggplot2 version 3.3.6.

### Statistics

Statistical tests are indicated in figure legends. Statistical analyses were performed using R software version 4.0.3 and GraphPad Prism software version 9.5.1. R packages used for statistical tests include rstatix version 0.7.2, ggpubr version 0.4.0, and stats version 4.0.3. Living/deceased status of transplant donor, HLA-A/B-DR mismatch, and induction immunosuppression were included as covariates in regression models in Supplementary Tables 1-2 were based on univariate analyses where *p* < 0.1. Additional covariates, including thymoglobulin induction, acute rejection, and delayed graft function, were added based on other studies that have shown correlations with outcome.

## DATA AVAILABILITY

Data from this study are available from corresponding authors upon request.

## ACKNOWLEDGEMENTS

The work was supported by NIH U01 AI63594 awarded to P.S.H with a supplement to A.H. D-F.R was funded by NIH T32 AI078892 Translational Immunology Training Grant. We thank the CTOT-01 and CTOT-19 patients and investigators. HLA genotyping was funded in whole or in part in M.C.’s lab with federal funds from the Frederick National Laboratory for Cancer Research, under contract no. 75N91019D00024, and was supported in part by the Intramural Research Program of the NIH, Frederick National Lab, Center for Cancer Research. Wildtype and HLA-E+ K562 cell lines were provided by Deepta Bhattacharya. Cell sorting was performed at the Dean’s Flow Cytometry CoRE at the Icahn School of Medicine at Mount Sinai and supported in part by the Tisch Cancer Institute at Mount Sinai P30 CA196521 – Cancer Center Support Grant. We thank the Human Immune Monitoring Center at Icahn School of Medicine at Mount Sinai for their support. The content of this publication does not necessarily reflect the views or policies of the Department of Health and Human Services, nor does mention of trade names, commercial products, or organizations imply endorsement by the U.S. Government.

## AUTHOR CONTRIBUTIONS

Oversight and funding by P.S.H and A.H. D-F.R performed and analyzed experiments, prepared figures, and wrote the initial draft of the manuscript. P.S.H and A.H edited and wrote the manuscript. M.F, N.C, and P.C helped with conceptualization of the project. Y.Y, M.M, and M.C provided support for HLA/KIR typing. M.M, G.K, B.L, R.M.R, R.R, G.G, and S.K.S helped with data acquisition.

