## Supplementary Tables for "Understanding the heterogeneity of alloreactive natural killer cell function in kidney transplantation"

**Supplementary Table 1A. Regression model of 2-year eGFR in CTOT01**

| | Estimate | P Val |
|---|---|---|
| Intercept | 57.9776 | 1e-04* |
| Thymoglobulin | 1.0779 | 0.8784 |
| Induction | -2.0563 | 0.8366 |
| **Living transplant** | **22.537** | **0.0024*** |
| Delayed Graft Function | 16.121 | 0.4734 |
| Acute Rejection | -1.8902 | 0.807 |
| HLA-A/B/C mismatch | -1.0399 | 0.5746 |
| **%Ksp37+** | **-4.8639** | **0.0408*** |

**Supplementary Table 1B. Regression model of 2-year eGFR in CTOT01**

| | Estimate | P Val |
|---|---|---|
| Intercept | 60.0487 | 2e-04* |
| Thymoglobulin | 1.1814 | 0.8665 |
| Induction | -2.5122 | 0.8018 |
| **Living transplant** | **21.8542** | **0.0041*** |
| Delayed Graft Function | 16.395 | 0.4649 |
| Acute Rejection | -1.7256 | 0.823 |
| HLA-A/B/DR mismatch | -1.3429 | 0.5145 |
| **%Ksp37+** | **-4.8235** | **0.0424*** |

**Supplementary Table 2A. Regression model of 5-year eGFR in CTOT01**

| | Estimate | P Val |
|---|---|---|
| Intercept | 66.9669 | 1e-04* |
| Thymoglobulin | -4.8275 | 0.5345 |
| Induction | -10.0663 | 0.3842 |
| Living transplant | 9.8193 | 0.2241 |
| Delayed Graft Function | 25.4289 | 0.3582 |
| Acute rejection | 4.7733 | 0.5894 |
| HLA-A/B/C mismatch | -0.4782 | 0.821 |
| **%Ksp37+** | **-5.992** | **0.0229*** |

**Supplementary Table 2B. Regression model of 5-year eGFR in CTOT01**

| | Estimate | P Val |
|---|---|---|
| Intercept | 69.9433 | 1e-04* |
| Thymoglobulin | -4.8539 | 0.5306 |
| Induction | -10.4846 | 0.3657 |
| Living transplant | 9.032 | 0.273 |
| Delayed Graft Function | 26.3281 | 0.3401 |
| Acute rejection | 4.8964 | 0.5792 |
| HLA-A/B/DR mismatch | -1.0378 | 0.6293 |
| **%Ksp37+** | **-5.961** | **0.0232*** |

**Supplementary Table 3A. Regression model of 6-month eGFR in CTOT19**

| | Estimate | P Val |
|---|---|---|
| Intercept | 54.749 | 0* |
| Delayed Graft Function | -0.443 | 0.9648 |
| **%Ksp37+** | **-1.825** | **0.0321*** |

**Supplementary Table 3B. Regression model of 6-month eGFR in CTOT19**

| | Estimate | P Val |
|---|---|---|
| Intercept | 64.2575 | 0* |
| HLA-A/B/C mismatch | -2.8126 | 0.1524 |
| %Ksp37+ | -1.533 | 0.0639 |

**Supplementary Table 3C. Regression model of 6-month eGFR in CTOT19**

| | Estimate | P Val |
|---|---|---|
| Intercept | 55.7924 | 0* |
| Acute Rejection | -5.4084 | 0.7523 |
| **%Ksp37+** | **-2.0835** | **0.0308*** |

**Supplementary Table 4A. Regression model of 2-year eGFR in CTOT19**

| | Estimate | P Val |
|---|---|---|
| Intercept | 64.5813 | 0* |
| Delayed Graft Function | -3.6054 | 0.7628 |
| **%Ksp37+** | **-2.011** | **0.0454*** |

**Supplementary Table 4B. Regression model of 2-year eGFR in CTOT19**

| | Estimate | P Val |
|---|---|---|
| Intercept | 70.9803 | 0* |
| HLA-A/B/C mismatch | -2.1225 | 0.3716 |
| %Ksp37+ | -1.7936 | 0.0761 |

**Supplementary Table 4C. Regression model of 2-year eGFR in CTOT19**

| | Estimate | P Val |
|---|---|---|
| Intercept | 64.0183 | 0* |
| Acute Rejection | -8.7277 | 0.6709 |
| %Ksp37+ | -2.13 | 0.0611 |

**Supplementary Table 5. Class I HLA alleles and KIR ligands in CTOT01**

| | HLA-A | | | HLA-B | | | | HLA-C | | |
|---|---|---|---|---|---|---|---|---|---|---|
| | Allele 1 | Allele 2 | Ligands | Allele 1 | Allele 2 | Ligands | -21 Dimorphism | Allele 1 | Allele 2 | Ligands |
| Donor #1 | 02:02 | 30:01 | NA | 07:02 | 53:01 | Bw4/Bw6 | MT | 04:01 | 07:02 | C1/C2 |
| Recipient #1 | 02:02 | 30:01 | NA | 42:01 | 53:01 | Bw4/Bw6 | MT | 04:01 | 17:01 | C2/C2 |
| Donor #2 | 01:01 | 32:01 | Bw4 | 08:01 | 40:01 | Bw6/Bw6 | MT | 02:02 | 07:01 | C1/C2 |
| Recipient #2 | 02:01 | 03:01 | A03 | 08:01 | 38:01 | Bw4/Bw6 | MM | 07:01 | 12:03 | C1/C1 |
| Donor #3 | 01:01 | 24:02 | Bw4 | 08:01 | 35:03 | Bw6/Bw6 | MT | 04:01 | 07:01 | C1/C2 |
| Recipient #3 | 01:01 | 24:02 | Bw4 | 08:01 | 35:03 | Bw6/Bw6 | MT | 04:01 | 07:01 | C1/C2 |
| Donor #4 | 02:01 | 02:01 | NA | 40:02 | 44:02 | Bw4/Bw6 | TT | 02:02 | 05:01 | C2/C2 |
| Recipient #4 | 01:01 | 02:01 | NA | 40:02 | 57:01 | Bw4/Bw6 | TT | 02:02 | 06:02 | C2/C2 |
| Donor #5 | 30:01 | 74:01 | NA | 08:01 | 44:03 | Bw4/Bw6 | MT | 07:01 | 14:03 | C1/C1 |
| Recipient #5 | 32:01 | 74:01 | Bw4 | 08:01 | 81:01 | Bw6/Bw6 | MM | 07:01 | 08:04 | C1/C1 |
| Donor #6 | 01:01 | 24:02 | Bw4 | 07:05 | 35:02 | Bw6/Bw6 | MT | 04:01 | 15:05 | C2/C2 |
| Recipient #6 | 23:01 | 24:02 | Bw4/Bw4 | 27:05 | 44:27 | Bw4/Bw4 | TT | 01:02 | 07:04 | C1/C1 |
| Donor #7 | 68:01 | 68:01 | NA | 15:15 | 35:01 | Bw6/Bw6 | TT | 01:02 | 07:02 | C1/C1 |
| Recipient #7 | 01:03 | 24:02 | Bw4 | 15:01 | 73:01 | Bw6/Bw6 | MT | 03:04 | 15:05 | C1/C2 |
| Donor #8 | 01:01 | 32:01 | Bw4 | 08:01 | 14:01 | Bw6/Bw6 | MM | 07:01 | 08:02 | C1/C1 |
| Recipient #8 | 02:01 | 03:01 | A03 | 35:01 | 51:01 | Bw4/Bw6 | TT | 04:01 | 15:02 | C2/C2 |
| Donor #9 | 02:01 | 32:01 | Bw4 | 07:02 | 14:01 | Bw6/Bw6 | MM | 07:02 | 08:02 | C1/C1 |
| Recipient #9 | 02:01 | 02:01 | NA | 07:02 | 08:01 | Bw6/Bw6 | MM | 07:01 | 07:02 | C1/C1 |
| Donor #10 | 02:05 | 31:01 | NA | 35:08 | 41:01 | Bw6/Bw6 | TT | 04:01 | 07:01 | C1/C2 |
| Recipient #10 | 23:01 | 66:01 | Bw4 | 08:01 | 58:02 | Bw4/Bw6 | MT | 03:04 | 06:02 | C1/C2 |
| Donor #11 | 01:01 | 03:01 | A03 | 07:02 | 08:01 | Bw6/Bw6 | MM | 07:01 | 07:02 | C1/C1 |
| Recipient #11 | 01:01 | 03:01 | A03 | 07:02 | 08:01 | Bw6/Bw6 | MM | 07:01 | 07:02 | C1/C1 |
| Donor #12 | 30:02 | 33:03 | NA | 15:16 | 57:03 | Bw4/Bw4 | TT | 07:01 | 14:02 | C1/C1 |
| Recipient #12 | 34:02 | 36:01 | NA | 35:01 | 53:01 | Bw4/Bw6 | TT | 04:01 | 04:01 | C2/C2 |
| Donor #13 | 23:01 | 33:03 | Bw4 | 15:16 | 53:01 | Bw4/Bw4 | TT | 06:02 | 14:02 | C1/C2 |
| Recipient #13 | 23:01 | 30:02 | Bw4 | 53:01 | 58:02 | Bw4/Bw4 | TT | 06:02 | 06:02 | C2/C2 |
| Donor #14 | 01:01 | 24:02 | Bw4 | 07:02 | 08:01 | Bw6/Bw6 | MM | 07:01 | 07:02 | C1/C1 |
| Recipient #14 | 01:01 | 03:01 | A03 | 08:01 | 15:18 | Bw6/Bw6 | MT | 07:01 | 07:04 | C1/C1 |
| Donor #15 | 02:01 | 66:01 | NA | 39:01 | 58:02 | Bw4/Bw6 | MT | 06:02 | 07:02 | C1/C2 |
| Recipient #15 | 02:01 | 33:03 | NA | 35:01 | 53:01 | Bw4/Bw6 | TT | 04:01 | 16:01 | C1/C2 |
| Donor #16 | 02:01 | 32:01 | Bw4 | 14:01 | 44:02 | Bw4/Bw6 | MT | 05:01 | 08:02 | C1/C2 |
| Recipient #16 | 02:01 | 03:01 | A03 | 40:01 | 40:02 | Bw6/Bw6 | TT | 02:02 | 03:04 | C1/C2 |
| Donor #17 | 26:01 | 32:01 | Bw4 | 41:02 | 44:02 | Bw4/Bw6 | TT | 05:01 | 17:03 | C2/C2 |
| Recipient #17 | 26:01 | 32:01 | Bw4 | 08:01 | 44:02 | Bw4/Bw6 | MT | 05:01 | 07:02 | C1/C2 |
| Donor #18 | 02:01 | 03:01 | A03 | 07:02 | 40:02 | Bw6/Bw6 | MT | 02:02 | 07:02 | C1/C2 |
| Recipient #18 | 02:01 | 25:01 | NA | 18:01 | 27:02 | Bw4/Bw6 | TT | 02:02 | 12:03 | C1/C2 |
| Donor #19 | 02:01 | 24:02 | Bw4 | 07:02 | 44:02 | Bw4/Bw6 | MT | 05:01 | 07:02 | C1/C2 |
| Recipient #19 | 01:01 | 24:02 | Bw4 | 27:05 | 44:02 | Bw4/Bw4 | TT | 02:02 | 05:01 | C2/C2 |
| Donor #20 | 02:01 | 24:02 | Bw4 | 40:01 | 40:02 | Bw6/Bw6 | TT | 03:04 | 03:04 | C1/C1 |
| Recipient #20 | 02:01 | 03:01 | A03 | 40:01 | 49:01 | Bw4/Bw6 | TT | 03:04 | 07:01 | C1/C1 |
| Donor #21 | 02:01 | 30:02 | NA | 13:02 | 18:01 | Bw4/Bw6 | TT | 05:01 | 06:02 | C2/C2 |
| Recipient #21 | 30:02 | 68:01 | NA | 08:01 | 18:01 | Bw6/Bw6 | MT | 05:01 | 07:01 | C1/C2 |
| Donor #22 | 24:02 | 31:01 | Bw4 | 51:01 | 51:01 | Bw4/Bw4 | TT | 05:01 | 14:02 | C1/C2 |
| Recipient #22 | 02:01 | 03:01 | A03 | 07:02 | 15:01 | Bw6/Bw6 | MT | 04:01 | 07:02 | C1/C2 |
| Donor #23 | 02:01 | 24:02 | Bw4 | 44:02 | 51:01 | Bw4/Bw4 | TT | 05:01 | 16:02 | C2/C2 |

| | | | | | | | | | | |
|---|---|---|---|---|---|---|---|---|---|---|
| Recipient #23 | 01:01 | 24:02 | Bw4 | 08:01 | 51:01 | Bw4/Bw6 | MT | 07:01 | 16:02 | C1/C2 |
| Donor #24 | 01:01 | 31:01 | NA | 51:01 | 52:01 | Bw4/Bw4 | TT | 02:02 | 12:02 | C1/C2 |
| Recipient #24 | 01:01 | 02:01 | NA | 44:02 | 52:01 | Bw4/Bw4 | TT | 05:01 | 12:02 | C1/C2 |
| Donor #25 | 01:01 | 01:01 | NA | 08:01 | 15:01 | Bw6/Bw6 | MT | 03:04 | 07:01 | C1/C1 |
| Recipient #25 | 01:01 | 03:01 | A03 | 08:01 | 53:01 | Bw4/Bw6 | MT | 04:01 | 07:01 | C1/C2 |
| Donor #26 | 24:02 | 32:01 | Bw4/Bw4 | 40:01 | 44:02 | Bw4/Bw6 | TT | 03:04 | 05:01 | C1/C2 |
| Recipient #26 | 24:02 | 25:01 | Bw4 | 39:01 | 44:02 | Bw4/Bw6 | MT | 05:01 | 12:03 | C1/C2 |
| Donor #27 | 01:01 | 03:01 | A03 | 07:02 | 08:01 | Bw6/Bw6 | MM | 07:01 | 07:02 | C1/C1 |
| Recipient #27 | 02:NA | 24:02 | Bw4 | 52:NA | 44:02 | Bw4 | TT | 05:01 | 07:02 | C1/C2 |
| Donor #28 | 30:01 | 30:01 | NA | 42:01 | 42:01 | Bw6/Bw6 | MM | 17:01 | 17:01 | C2/C2 |
| Recipient #28 | 03:01 | 33:03 | A03 | 07:02 | 53:01 | Bw4/Bw6 | MT | 04:01 | 07:02 | C1/C2 |
| Donor #29 | 02:01 | 02:01 | NA | 14:01 | 44:02 | Bw4/Bw6 | MT | 05:01 | 08:02 | C1/C2 |
| Recipient #29 | 29:02 | 36:01 | NA | 45:01 | 53:01 | Bw4/Bw6 | TT | 04:01 | 16:01 | C1/C2 |
| Donor #30 | 30:01 | 30:02 | NA | 42:01 | 58:01 | Bw4/Bw6 | MT | 07:01 | 17:01 | C1/C2 |
| Recipient #30 | 02:01 | 30:02 | NA | 07:02 | 58:01 | Bw4/Bw6 | MT | 07:01 | 07:02 | C1/C1 |
| Donor #31 | 01:01 | 32:01 | Bw4 | 08:01 | 55:01 | Bw6/Bw6 | MT | 05:01 | 07:01 | C1/C2 |
| Recipient #31 | 01:01 | 02:01 | NA | 08:01 | 15:03 | Bw6/Bw6 | MT | 02:10 | 07:01 | C1/C2 |
| Donor #32 | 02:01 | 32:01 | Bw4 | 15:01 | 44:02 | Bw4/Bw6 | TT | 03:03 | 05:01 | C1/C2 |
| Recipient #32 | 02:01 | 32:01 | Bw4 | 15:01 | 44:02 | Bw4/Bw6 | TT | 03:03 | 05:01 | C1/C2 |
| Donor #33 | 02:01 | 02:01 | NA | 08:01 | 40:01 | Bw6/Bw6 | MT | 03:04 | 07:01 | C1/C1 |
| Recipient #33 | 02:01 | 29:02 | NA | 40:01 | 44:03 | Bw4/Bw6 | TT | 03:04 | 16:01 | C1/C1 |
| Donor #34 | 02:01 | 31:01 | NA | 44:02 | 51:01 | Bw4/Bw4 | TT | 05:01 | 15:02 | C2/C2 |
| Recipient #34 | 02:01 | 31:01 | NA | 48:01 | 51:01 | Bw4/Bw6 | MT | 08:01 | 15:02 | C1/C2 |
| Donor #35 | 02:01 | 68:01 | NA | 35:01 | 44:02 | Bw4/Bw6 | TT | 05:01 | 15:02 | C2/C2 |
| Recipient #35 | 29:02 | 68:01 | NA | 35:01 | 44:03 | Bw4/Bw6 | TT | 15:02 | 16:01 | C1/C2 |
| Donor #36 | 02:01 | 32:01 | Bw4 | 15:01 | 39:01 | Bw6/Bw6 | MT | 03:04 | 07:02 | C1/C1 |
| Recipient #36 | 01:01 | 32:01 | Bw4 | 07:02 | 39:01 | Bw6/Bw6 | MM | 07:02 | 07:02 | C1/C1 |
| Donor #37 | 01:01 | 11:01 | A11 | 49:01 | 51:01 | Bw4/Bw4 | TT | 07:01 | 15:02 | C1/C2 |
| Recipient #37 | 02:01 | 03:01 | A03 | 07:02 | 15:01 | Bw6/Bw6 | MT | 03:04 | 07:02 | C1/C1 |
| Donor #38 | 02:01 | 02:04 | NA | 40:01 | 51:01 | Bw4/Bw6 | TT | 03:04 | 15:02 | C1/C2 |
| Recipient #38 | 24:02 | 24:02 | Bw4/Bw4 | 07:02 | 44:03 | Bw4/Bw6 | MT | 04:01 | 07:02 | C1/C2 |
| Donor #39 | 03:01 | 29:02 | A03 | 14:02 | 44:03 | Bw4/Bw6 | MT | 08:02 | 16:01 | C1/C1 |
| Recipient #39 | 02:01 | 24:02 | Bw4 | 40:01 | 45:01 | Bw6/Bw6 | TT | 03:04 | 16:01 | C1/C1 |
| Donor #40 | 01:01 | 11:01 | A11 | 08:01 | 56:01 | Bw6/Bw6 | MT | 01:02 | 07:01 | C1/C1 |
| Recipient #40 | 02:01 | 29:01 | NA | 18:01 | 57:01 | Bw4/Bw6 | TT | 06:02 | 07:01 | C1/C2 |
| Donor #41 | 01:01 | 32:01 | Bw4 | 13:02 | 40:02 | Bw4/Bw6 | TT | 02:02 | 06:02 | C2/C2 |
| Recipient #41 | 01:01 | 02:01 | NA | 08:01 | 15:01 | Bw6/Bw6 | MT | 03:03 | 07:01 | C1/C1 |
| Donor #42 | 23:01 | 34:02 | Bw4 | 15:03 | 52:01 | Bw4/Bw6 | TT | 02:10 | 16:01 | C1/C2 |
| Recipient #42 | 11:01 | 29:02 | A11 | 07:02 | 07:02 | Bw6/Bw6 | MM | 07:02 | 07:02 | C1/C1 |
| Donor #43 | 34:02 | 74:01 | NA | 44:03 | 57:03 | Bw4/Bw4 | TT | 04:01 | 07:01 | C1/C2 |
| Recipient #43 | 02:02 | 34:02 | NA | 15:16 | 44:03 | Bw4/Bw4 | TT | 04:01 | 14:02 | C1/C2 |
| Donor #44 | 01:01 | 23:17 | Bw4 | 08:01 | 15:03 | Bw6/Bw6 | MT | 02:10 | 07:02 | C1/C2 |
| Recipient #44 | 25:01 | 30:01 | NA | 07:02 | 53:01 | Bw4/Bw6 | MT | 04:01 | 07:02 | C1/C2 |
| Donor #45 | 03:01 | 68:02 | A03 | 15:10 | 15:10 | Bw6/Bw6 | TT | 03:04 | 03:04 | C1/C1 |
| Recipient #45 | 23:01 | 68:02 | Bw4 | 07:02 | 15:10 | Bw6/Bw6 | MT | 03:04 | 15:05 | C1/C2 |
| Donor #46 | 02:03 | 26:01 | NA | 39:09 | 52:01 | Bw4/Bw6 | MT | 07:02 | 07:02 | C1/C1 |
| Recipient #46 | 68:02 | 68:02 | NA | 15:10 | 35:01 | Bw6/Bw6 | TT | 03:04 | 04:01 | C1/C2 |
| Donor #47 | 03:01 | 29:02 | A03 | 07:02 | 44:03 | Bw4/Bw6 | MT | 07:02 | 16:01 | C1/C1 |

| | | | | | | | | | | | |
|---|---|---|---|---|---|---|---|---|---|---|---|
| Recipient #47 | 03:01 | 29:02 | A03 | 07:02 | 44:03 | Bw4/Bw6 | MT | | 04:01 | 07:02 | C1/C2 |
| Donor #48 | 02:01 | 03:01 | A03 | 14:02 | 51:01 | Bw4/Bw6 | MT | | 08:02 | 12:03 | C1/C1 |
| Recipient #48 | 03:01 | 26:01 | A03 | 15:01 | 38:01 | Bw4/Bw6 | MT | | 03:03 | 12:03 | C1/C1 |
| Donor #49 | 02:01 | 03:01 | A03 | 14:02 | 51:01 | Bw4/Bw6 | MT | | 08:02 | 12:03 | C1/C1 |
| Recipient #49 | 11:01 | 23:01 | A11/Bw4 | 57:01 | 58:01 | Bw4/Bw4 | TT | | 06:02 | 07:18 | C1/C2 |
| Donor #50 | 03:01 | 68:02 | A03 | 14:02 | 57:01 | Bw4/Bw6 | MT | | 06:02 | 08:02 | C1/C2 |
| Recipient #50 | 02:01 | 11:01 | A11 | 18:01 | 35:01 | Bw6/Bw6 | TT | | 04:01 | 07:01 | C1/C2 |
| Donor #51 | 33:03 | 68:01 | NA | 51:01 | 58:02 | Bw4/Bw4 | TT | | 06:02 | 16:01 | C1/C2 |
| Recipient #51 | 29:02 | 66:01 | NA | 44:03 | 58:02 | Bw4/Bw4 | TT | | 04:01 | 06:02 | C2/C2 |
| Donor #52 | 23:01 | 32:01 | Bw4/Bw4 | 13:02 | 50:01 | Bw4/Bw6 | TT | | 06:02 | 06:02 | C2/C2 |
| Recipient #52 | 02:01 | 23:01 | Bw4 | 27:05 | 50:01 | Bw4/Bw6 | TT | | 01:02 | 06:02 | C1/C2 |
| Donor #53 | 01:01 | 25:01 | NA | 08:01 | 58:01 | Bw4/Bw6 | MT | | 07:01 | 07:18 | C1/C1 |
| Recipient #53 | 01:01 | 25:01 | NA | 08:01 | 58:01 | Bw4/Bw6 | MT | | 07:01 | 07:18 | C1/C1 |
| Donor #54 | 02:01 | 24:02 | Bw4 | 15:01 | 44:02 | Bw4/Bw6 | TT | | 03:03 | 05:01 | C1/C2 |
| Recipient #54 | 29:02 | 66:01 | NA | 47:03 | 53:01 | Bw4/Bw6 | TT | | 04:01 | 07:01 | C1/C2 |
| Donor #55 | 02:01 | 68:01 | NA | 35:01 | 35:08 | Bw6/Bw6 | TT | | 02:02 | 04:01 | C2/C2 |
| Recipient #55 | 30:01 | 74:01 | NA | 35:01 | 53:01 | Bw4/Bw6 | TT | | 04:01 | 04:01 | C2/C2 |
| Donor #56 | 01:01 | 02:01 | NA | 08:01 | 44:02 | Bw4/Bw6 | MT | | 05:01 | 07:01 | C1/C2 |
| Recipient #56 | 03:01 | 11:01 | A03/A11 | 35:01 | 53:01 | Bw4/Bw6 | TT | | 04:01 | 04:01 | C2/C2 |
| Donor #57 | 30:02 | 68:01 | NA | 49:01 | 51:01 | Bw4/Bw4 | TT | | 07:01 | 15:02 | C1/C2 |
| Recipient #57 | 02:01 | 29:02 | NA | 07:06 | 35:01 | Bw6/Bw6 | MT | | 04:01 | 07:02 | C1/C2 |
| Donor #58 | 02:01 | 02:01 | NA | 15:01 | 18:01 | Bw6/Bw6 | TT | | 03:04 | 12:03 | C1/C1 |
| Recipient #58 | 02:01 | 33:03 | NA | 07:02 | 53:01 | Bw4/Bw6 | MT | | 04:01 | 07:02 | C1/C2 |
| Donor #59 | 02:05 | 23:01 | Bw4 | 44:03 | 50:01 | Bw4/Bw6 | TT | | 04:01 | 06:02 | C2/C2 |
| Recipient #59 | 03:01 | 23:01 | A03/Bw4 | 07:02 | 44:03 | Bw4/Bw6 | MT | | 04:01 | 07:02 | C1/C2 |
| Donor #60 | 02:01 | 23:01 | Bw4 | 49:01 | 51:01 | Bw4/Bw4 | TT | | 03:03 | 07:01 | C1/C1 |
| Recipient #60 | 02:01 | 26:01 | NA | 38:01 | 55:01 | Bw4/Bw6 | MT | | 03:03 | 12:03 | C1/C1 |
| Donor #61 | 01:01 | 03:01 | A03 | 35:01 | 57:01 | Bw4/Bw6 | TT | | 04:01 | 06:02 | C2/C2 |
| Recipient #61 | 01:01 | 03:01 | A03 | 35:01 | 57:01 | Bw4/Bw6 | TT | | 04:01 | 06:02 | C2/C2 |
| Donor #62 | 24:02 | 32:01 | Bw4/Bw4 | 35:01 | 57:01 | Bw4/Bw6 | TT | | 03:03 | 06:02 | C1/C2 |
| Recipient #62 | 03:01 | 23:01 | A03/Bw4 | 15:03 | 42:01 | Bw6/Bw6 | MT | | 02:10 | 17:01 | C2/C2 |
| Donor #63 | 11:01 | 33:03 | A11 | 07:02 | 18:01 | Bw6/Bw6 | MT | | 05:01 | 15:05 | C2/C2 |
| Recipient #63 | 11:01 | 32:01 | A11/Bw4 | 08:01 | 18:01 | Bw6/Bw6 | MT | | 05:01 | 07:01 | C1/C2 |
| Donor #64 | 36:01 | 68:01 | NA | 52:01 | 53:01 | Bw4/Bw4 | TT | | 04:01 | 16:01 | C1/C2 |
| Recipient #64 | 02:02 | 68:01 | NA | 15:03 | 52:01 | Bw4/Bw6 | TT | | 02:10 | 16:01 | C1/C2 |
| Donor #65 | 02:01 | 68:02 | NA | 14:02 | 44:02 | Bw4/Bw6 | MT | | 05:09 | 08:02 | C1/C2 |
| Recipient #65 | 02:01 | 30:01 | NA | 15:03 | 40:02 | Bw6/Bw6 | TT | | 02:10 | 15:02 | C2/C2 |
| Donor #66 | 68:01 | 68:02 | NA | 45:01 | 49:01 | Bw4/Bw6 | TT | | 06:02 | 07:01 | C1/C2 |
| Recipient #66 | 02:01 | 24:02 | Bw4 | 35:01 | 35:12 | Bw6/Bw6 | TT | | 04:01 | 04:01 | C2/C2 |
| Donor #67 | 01:01 | 74:01 | NA | 08:01 | 35:01 | Bw6/Bw6 | MT | | 07:01 | 07:01 | C1/C1 |
| Recipient #67 | 01:01 | 33:03 | NA | 18:01 | 57:03 | Bw4/Bw6 | TT | | 04:01 | 07:01 | C1/C2 |
| Donor #68 | 02:05 | 23:01 | Bw4 | 07:02 | 35:01 | Bw6/Bw6 | MT | | 04:01 | 07:01 | C1/C2 |
| Recipient #68 | 23:01 | 30:02 | Bw4 | 15:10 | 35:01 | Bw6/Bw6 | TT | | 03:04 | 07:01 | C1/C1 |
| Donor #69 | 03:01 | 30:02 | A03 | 14:03 | 42:02 | Bw6/Bw6 | MM | | 08:02 | 17:01 | C1/C2 |
| Recipient #69 | 30:02 | 33:03 | NA | 42:02 | 58:01 | Bw4/Bw6 | MT | | 03:02 | 17:01 | C1/C2 |
| Donor #70 | 03:01 | 74:01 | A03 | 07:02 | 35:01 | Bw6/Bw6 | MT | | 07:01 | 07:02 | C1/C1 |
| Recipient #70 | 02:01 | 03:01 | A03 | 35:01 | 45:01 | Bw6/Bw6 | TT | | 07:01 | 16:01 | C1/C1 |

**Supplementary Table 6. Class I HLA allele and KIR ligands in CTOT19**

| | HLA-A | | | HLA-B | | | | HLA-C | | |
|---|---|---|---|---|---|---|---|---|---|---|
| ID | Allele 1 | Allele 2 | Ligands | Allele 1 | Allele 2 | Ligands | -21 Dimorphism | Allele 1 | Allele 2 | Ligands |
| Donor #1 | 02:01 | 25:01 | NA | 55:01 | 58:01 | Bw4/Bw6 | TT | 03:03 | 07:18 | C1/C1 |
| Recipient #1 | 74:01 | 74:01 | NA | 27:05 | 35:01 | Bw4/Bw6 | TT | 02:10 | 04:01 | C2/C2 |
| Donor #2 | 01:01 | 03:01 | A03 | 35:01 | 44:02 | Bw4/Bw6 | TT | 04:01 | 05:01 | C2/C2 |
| Recipient #2 | 03:01 | 68:01 | A03 | 07:02 | 44:02 | Bw4/Bw6 | MT | 05:01 | 07:02 | C1/C2 |
| Donor #3 | 01:01 | 02:01 | NA | 15:01 | 50:02 | Bw6/Bw6 | TT | 03:03 | 06:02 | C1/C2 |
| Recipient #3 | 01:01 | 02:01 | NA | 15:01 | 50:02 | Bw6/Bw6 | TT | 03:03 | 06:02 | C1/C2 |
| Donor #4 | 26:01 | 30:02 | NA | 07:02 | 51:01 | Bw4/Bw6 | MT | 07:02 | 14:02 | C1/C1 |
| Recipient #4 | 03:01 | 30:02 | A03 | 07:02 | 57:03 | Bw4/Bw6 | MT | 07:02 | 07:18 | C1/C1 |
| Donor #5 | 02:01 | 24:02 | Bw4 | 07:02 | 15:01 | Bw6/Bw6 | MT | 03:04 | 07:02 | C1/C1 |
| Recipient #5 | 02:01 | 24:02 | Bw4 | 07:02 | 15:01 | Bw6/Bw6 | MT | 03:04 | 07:02 | C1/C1 |
| Donor #6 | 24:02 | 30:01 | Bw4 | 13:02 | 39:06 | Bw4/Bw6 | MT | 06:02 | 07:02 | C1/C2 |
| Recipient #6 | 30:01 | 33:03 | NA | 42:01 | 53:01 | Bw4/Bw6 | MT | 04:01 | 17:01 | C2/C2 |
| Donor #7 | 02:01 | 68:02 | NA | 15:10 | 44:02 | Bw4/Bw6 | TT | 03:04 | 05:01 | C1/C2 |
| Recipient #7 | 02:01 | 68:01 | NA | 15:18 | 44:02 | Bw4/Bw6 | TT | 05:01 | 07:04 | C1/C2 |
| Donor #8 | 24:02 | 30:01 | Bw4 | 18:01 | 42:01 | Bw6/Bw6 | MT | 07:01 | 17:01 | C1/C2 |
| Recipient #8 | 30:01 | 30:01 | NA | 15:03 | 42:01 | Bw6/Bw6 | MT | 02:10 | 17:01 | C2/C2 |
| Donor #9 | 02:01 | 26:01 | NA | 18:01 | 38:01 | Bw4/Bw6 | MT | 07:01 | 12:03 | C1/C1 |
| Recipient #9 | 03:01 | 68:02 | A03 | 42:01 | 44:02 | Bw4/Bw6 | MT | 07:04 | 17:01 | C1/C2 |
| Donor #10 | 01:01 | 26:01 | NA | 27:05 | 44:03 | Bw4/Bw4 | TT | 01:02 | 04:01 | C1/C2 |
| Recipient #10 | 01:01 | 26:01 | NA | 27:05 | 44:03 | Bw4/Bw4 | TT | 01:02 | 04:01 | C1/C2 |
| Donor #11 | 02:01 | 02:01 | NA | 44:02 | 51:01 | Bw4/Bw4 | TT | 05:01 | 15:02 | C2/C2 |
| Recipient #11 | 02:01 | 24:02 | Bw4 | 15:01 | 35:01 | Bw6/Bw6 | TT | 04:01 | 04:01 | C2/C2 |
| Donor #12 | 11:01 | 31:01 | A11 | 40:02 | 44:03 | Bw4/Bw6 | TT | 03:04 | 16:01 | C1/C1 |
| Recipient #12 | 02:01 | 66:01 | NA | 15:15 | 41:02 | Bw6/Bw6 | TT | 01:02 | 17:03 | C1/C2 |
| Donor #13 | 03:01 | 68:01 | A03 | 18:01 | 51:01 | Bw4/Bw6 | TT | 05:01 | 15:02 | C2/C2 |
| Recipient #13 | 03:01 | 68:01 | A03 | 18:01 | 51:01 | Bw4/Bw6 | TT | 05:01 | 15:02 | C2/C2 |
| Donor #14 | 11:01 | 24:02 | A11/Bw4 | 15:01 | 18:01 | Bw6/Bw6 | TT | 03:03 | 12:03 | C1/C1 |
| Recipient #14 | 30:01 | 30:02 | NA | 15:03 | 42:01 | Bw6/Bw6 | MT | 02:10 | 17:01 | C2/C2 |
| Donor #15 | 23:01 | 31:01 | Bw4 | 44:03 | 55:01 | Bw4/Bw6 | TT | 03:03 | 04:09 | C1/C2 |
| Recipient #15 | 01:01 | 03:01 | A03 | 07:02 | 40:01 | Bw6/Bw6 | MT | 03:04 | 07:02 | C1/C1 |
| Donor #16 | 23:01 | 31:01 | Bw4 | 44:03 | 55:01 | Bw4/Bw6 | TT | 03:03 | 04:09 | C1/C2 |
| Recipient #16 | 01:01 | 02:01 | NA | 08:01 | 49:01 | Bw4/Bw6 | MT | 07:01 | 07:01 | C1/C1 |
| Donor #17 | 24:02 | 29:02 | Bw4 | 13:02 | 55:01 | Bw4/Bw6 | TT | 01:02 | 06:02 | C1/C2 |
| Recipient #17 | 02:01 | 23:01 | Bw4 | 07:02 | 41:01 | Bw6/Bw6 | MT | 07:01 | 07:02 | C1/C1 |
| Donor #18 | 01:01 | 32:01 | Bw4 | 15:01 | 51:01 | Bw4/Bw6 | TT | 03:03 | 14:02 | C1/C1 |
| Recipient #18 | 11:01 | 31:01 | A11 | 44:02 | 51:01 | Bw4/Bw4 | TT | 05:01 | 15:02 | C2/C2 |
| Donor #19 | 02:01 | 11:01 | A11 | 51:01 | 56:01 | Bw4/Bw6 | TT | 01:02 | 02:02 | C1/C2 |
| Recipient #19 | 23:17 | 68:01 | Bw4 | 08:01 | 39:05 | Bw6/Bw6 | MM | 07:01 | 07:02 | C1/C1 |
| Donor #20 | 11:01 | 24:02 | A11/Bw4 | 13:02 | 35:01 | Bw4/Bw6 | TT | 04:01 | 06:02 | C2/C2 |
| Recipient #20 | 03:01 | 03:01 | A03/A03 | 35:01 | 45:01 | Bw6/Bw6 | TT | 04:01 | 05:01 | C2/C2 |
| Donor #21 | 01:01 | 24:02 | Bw4 | 07:02 | 35:01 | Bw6/Bw6 | MT | 04:01 | 07:02 | C1/C2 |
| Recipient #21 | 02:02 | 30:01 | NA | 42:02 | 53:01 | Bw4/Bw6 | MT | 04:01 | 17:01 | C2/C2 |

| | | | | | | | | | | |
|---|---|---|---|---|---|---|---|---|---|---|
| Donor #22 | 02:01 | 11:01 | A11 | 15:01 | 51:01 | Bw4/Bw6 | TT | 03:03 | 15:02 | C1/C2 |
| Recipient #22 | 02:01 | 03:01 | A03 | 07:02 | 51:01 | Bw4/Bw6 | MT | 07:02 | 14:02 | C1/C1 |
| Donor #23 | 24:02 | 26:01 | Bw4 | 38:01 | 48:01 | Bw4/Bw6 | MM | 08:03 | 12:03 | C1/C1 |
| Recipient #23 | 02:01 | 33:03 | NA | 45:01 | 58:01 | Bw4/Bw6 | TT | 03:02 | 16:01 | C1/C1 |
| Donor #24 | 03:01 | 30:02 | A03 | 15:03 | 35:01 | Bw6/Bw6 | TT | 02:10 | 04:01 | C2/C2 |
| Recipient #24 | 66:01 | 68:02 | NA | 35:01 | 41:02 | Bw6/Bw6 | TT | 16:01 | 17:03 | C1/C2 |
| Donor #25 | 02:01 | 68:03 | NA | 39:05 | 52:01 | Bw4/Bw6 | MT | 03:03 | 07:02 | C1/C1 |
| Recipient #25 | 24:02 | 26:01 | Bw4 | 35:14 | 57:01 | Bw4/Bw6 | TT | 04:01 | 06:02 | C2/C2 |
| Donor #26 | 02:01 | 02:01 | NA | 44:02 | 45:01 | Bw4/Bw6 | TT | 05:01 | 06:02 | C2/C2 |
| Recipient #26 | 02:01 | 02:01 | NA | 44:02 | 45:01 | Bw4/Bw6 | TT | 05:01 | 06:02 | C2/C2 |

**Supplementary Table 7. Class I HLA allele and KIR ligands in healthy cohort**

| | HLA-A | | | HLA-B | | | | HLA-C | | |
|---|---|---|---|---|---|---|---|---|---|---|
| ID | Allele 1 | Allele 2 | Ligands | Allele 1 | Allele 2 | Ligands | -21 Dimorphism | Allele 1 | Allele 2 | Ligands |
| Healthy #1 | 31:01 | 32:01 | Bw4 | 15:01 | 40:01 | Bw6/Bw6 | TT | 03:03 | 03:04 | C1/C1 |
| Healthy #2 | 24:02 | 30:02 | Bw4 | 18:01 | 35:01 | Bw6/Bw6 | TT | 04:01 | 06:02 | C2/C2 |
| Healthy #3 | 23:01 | 31:01 | Bw4 | 38:01 | 49:01 | Bw4/Bw4 | MT | 07:01 | 12:03 | C1/C1 |
| Healthy #4 | 02:01 | 11:01 | A11 | 51:01 | 52:01 | Bw4/Bw4 | TT | 12:02 | 15:02 | C1/C2 |
| Healthy #5 | 24:02 | 66:01 | Bw4 | 38:01 | 52:01 | Bw4/Bw4 | MT | 12:02 | 12:03 | C1/C1 |
| Healthy #6 | 24:02 | 33:03 | Bw4 | 13:01 | 53:01 | Bw4/Bw4 | TT | 03:04 | 04:01 | C1/C2 |
| Healthy #7 | 25:01 | 31:01 | NA | 18:01 | 35:12 | Bw6/Bw6 | TT | 04:01 | 12:03 | C1/C2 |
| Healthy #8 | 02:05 | 26:01 | NA | 41:01 | 52:01 | Bw4/Bw6 | TT | 07:01 | 12:02 | C1/C1 |
| Healthy #9 | 23:01 | 24:02 | Bw4/Bw4 | 18:01 | 18:01 | Bw6/Bw6 | TT | 07:01 | 07:04 | C1/C1 |
| Healthy #10 | 02:01 | 03:01 | A03 | 35:01 | 51:01 | Bw4/Bw6 | TT | 04:01 | 15:02 | C2/C2 |
| Healthy #11 | 02:01 | 30:04 | NA | 44:02 | 49:01 | Bw4/Bw4 | TT | 05:01 | 07:01 | C1/C2 |
| Healthy #12 | 32:01 | 33:01 | Bw4 | 08:01 | 14:02 | Bw6/Bw6 | MM | 07:01 | 08:02 | C1/C1 |
| Healthy #13 | 24:02 | 26:01 | Bw4 | 18:01 | 38:01 | Bw4/Bw6 | MT | 12:03 | 12:03 | C1/C1 |
| Healthy #14 | 01:01 | 02:01 | NA | 08:01 | 57:01 | Bw4/Bw6 | MT | 06:02 | 07:01 | C1/C2 |
| Healthy #15 | 30:02 | 68:01 | NA | 15:03 | 53:01 | Bw4/Bw6 | TT | 02:10 | 04:01 | C2/C2 |
| Healthy #16 | 02:01 | 26:01 | NA | 08:01 | 53:01 | Bw4/Bw6 | MT | 03:04 | 04:01 | C1/C2 |
| Healthy #17 | 11:01 | 24:02 | A11/Bw4 | 15:01 | 15:25 | Bw6/Bw6 | TT | 03:03 | 07:26 | C1/C1 |
| Healthy #18 | 02:01 | 03:01 | A03 | 51:01 | 53:01 | Bw4/Bw4 | TT | 01:02 | 16:01 | C1/C1 |
| Healthy #19 | 01:01 | 02:03 | NA | 40:06 | 57:01 | Bw4/Bw6 | TT | 06:02 | 15:02 | C2/C2 |
| Healthy #20 | 01:01 | 26:01 | NA | 44:02 | 53:01 | Bw4/Bw4 | TT | 04:01 | 05:01 | C2/C2 |

**Supplementary Table 8. Flow cytometry panel for profiling stimulator cell ligands**

| | Antigen | Vendor | Catalog # | Clone |
|---|---|---|---|---|
| Panel 1 | HLA-A/B/C | BioLegend | 311418 | W6/32 |
| | MICA/B | BioLegend | 320912 | 6D4 |
| | CD86 | BioLegend | 374214 | BU63 |
| | HLA-F | BioLegend | 373208 | 3D11/HLA-F |
| | CD112 | Miltenyi | 130-122-770 | REA1195 |
| Panel 2 | CD80 | BioLegend | 305225 | 2D10 |
| | HLA-G | BioLegend | 335912 | 87G |
| | ULBP1 | R&D Systems | FAB1380A | 170818 |
| | CD54 | BioLegend | 353117 | HA58 |
| | HLA-C | BD Biosciences | 747594 | DT-9 |
| Panel 3 | CD155 | BioLegend | 337634 | SKII.4 |
| | HLA-E | BioLegend | 342604 | 3D12 |
| | PD-L1 | BioLegend | 329736 | 29E.2A3 |
| | ULBP2/5/6 | R&D Systems | FAB1298G | 165903 |
| | CD58 | BioLegend | 330918 | TS2/9 |
| Viability | Zombie NIR | BioLegend | 423105 | |

**Supplementary Table 9. CyTOF Panel**

| Metal Conjugate | Antigen | Vendor | Catalog # | Clone | Function |
|---|---|---|---|---|---|
| 89Y | CD45 | Fluidigm | 3089003B | HI30 | Lineage marker; leukocyte |
| 111Cd | LILRB1 | BioLegend | 333702 | GHI/75 | NK cell |
| 112Cd | CD8 | BioLegend | 301002 | RPA-T8 | Lineage marker; T cell |
| 113Cd | CD38 | Miltenyi | 130-122-307 | REA572 | NK cell, B cell |
| 114Cd | CD3 | BioLegend | 300402 | UCHT1 | Lineage marker; T cell |
| 115In | CD14 | BioLegend | 301802 & 301843 | M5E2 | Lineage marker; myeloid & other |
| 115In | CD19 | BioLegend | 302202 | HIB19 | Lineage marker; B cell |
| 115In | CD33 | BioLegend | 303402 | WM53 | Lineage marker; myeloid & other |
| 116Cd | CD57 | BioLegend | 359602 | HNK-1 | NK and T cell terminal maturation |
| 141Pr | TCRvd2 | BioLegend | 331402 | B6 | T cell |
| 142Nd | CCL4 | Miltenyi | 130-095-212 | REA511 | NK cell effector function |
| 143Nd | XCL1 | R&D Sys | AF695 | 109001 | NK cell effector function |
| 144Nd | Ksp37 | BioLegend | custom | TDA3 | NK cell effector function |
| 145Nd | NKG2D | Miltenyi | 130-122-332 | REA797 | NK cell |
| 146Nd | DNAM-1 | Miltenyi | 130-092-479 & 130-126-485 | DX11 & REA1040 | NK cell |
| 147Sm | PLZF | R&D Sys | MAB2944 | 6318100 | Transcription factor |
| 148Nd | KIR3DL1/L2 | Miltenyi | 130-126-489 | REA970 | NK cell education |
| 149Sm | CD25 | Fluidigm | 3149010B | 2A3 | NK and T cell effector function |
| 150Nd | TCF-1/7 | BioLegend | 655202 | 7F11A10 | Transcription factor |
| 151Eu | CCL5 | R&D Sys | MAB278-100 | 21445 | NK cell effector function |
| 152Sm | KIR3DL1 | Miltenyi | 130-092-555 | DX9 | NK cell education |
| 153Eu | TIM-3 | Fluidigm | 3153008B | F38-2E2 | Exhaustion |
| 154Sm | TIGIT | Fluidigm | 3154016B | MBSA43 | Exhaustion |
| 155Gd | NKp46 | Miltenyi | 130-124-522 | REA808 | NK cell |
| 156Gd | KIR2DL2/L3 | Miltenyi | 130-122-346 | REA1006 | NK cell education |
| 158Gd | KIR2DL1 | Miltenyi | 130-122-279 | REA284 | NK cell education |
| 159Tb | CD56 | Miltenyi | 130-108-016 | REA196 | NK cell |
| 160Gd | NKG2A | Miltenyi | 130-122-329 | REA110 | NK cell education |
| 161Dy | Ki-67 | Fluidigm | 3161007B | B56 | Proliferation |
| 162Dy | CD27 | Fluidigm | 3162009B | L128 | NK cell, T cell, B cell |
| 163Dy | CXCR3 | Fluidigm | 3163004B | G025H7 | NK and T cell effector function |
| 164Dy | NKG2C | Miltenyi | 130-122-278 | REA205 | NK cell |
| 165Ho | KIR2DL3 | Miltenyi | 130-122-280 | REA147 | NK cell education |
| 166Er | KIR2DL1/S1 | Miltenyi | 130-122-345 | REA1010 | NK cell education |
| 167Er | FceR1g | EMD millipore | 06-727 | polyclonal | NK cell education |
| 168Er | IFNg | Fluidigm | 3168005B | B27 | NK cell effector function |
| 169Tm | Granzyme K | BioLegend | 370502 | GM26E7 | NK cell effector function |
| 170Er | CD122 | Fluidigm | 3170004B | Tu27 | NK and T cell effector function |
| 171Yb | Granzyme B | Fluidigm | 3171002B | GB11 | NK cell effector function |
| 172Yb | CD107a | Miltenyi | 130-124-536 | REA792 | NK cell effector function |
| 173Yb | CD137 | Fluidigm | 3173015B | 4B4-1 | NK cell effector function |
| 174Yb | TOX | Miltenyi | custom | REA473 | Transcription factor |
| 175Lu | Perforin | Fluidigm | 3175004B | B-D48 | NK cell effector function |
| 176Yb | CD4 | Fluidigm | 3176010B | RPA-T4 | T cell |
| 198Pt | HLA-DR | Miltenyi | 130-122-299 | REA805 | Lineage marker; myeloid & other |
| 209Bi | CD16 | Fluidigm | 3209002B | 3G8 | NK cell |

**Supplementary Table 10. Flow cytometry panel for NK cell sorting and killing assay**

| Antigen | Vendor | Catalog # | Clone | Function |
|---|---|---|---|---|
| CD19 | BioLegend | 302244 | HIB19 | B cell |
| CD56 | BioLegend | 362550 | 5.1H11 | NK cell |
| CD3 | BioLegend | 317346 | OKT3 | T cell |
| NKG2A | Miltenyi | 130-128-163 | REA110 | NK cell education |
| NKG2C | Miltenyi | 130-117-398 | REA205 | NK cell education |
| KIR3DL1/L2 | Miltenyi | 130-116-180 | REA970 | NK cell education |
| KIR2D | Miltenyi | 130-117-483 | REA1042 | NK cell education |
| CD107a | Miltenyi | 130-111-624 | REA792 | NK cell effector function |
| Ksp37 | BioLegend | 346603 | TDA3 | NK cell effector function |
| Zombie NIR | BioLegend | 423106 | | Fixable viability dye |
| Propidium iodide | Life Technologies | P3566 | | Viability dye |
