## Supplementary Figures for "Understanding the heterogeneity of alloreactive natural killer cell function in kidney transplantation"

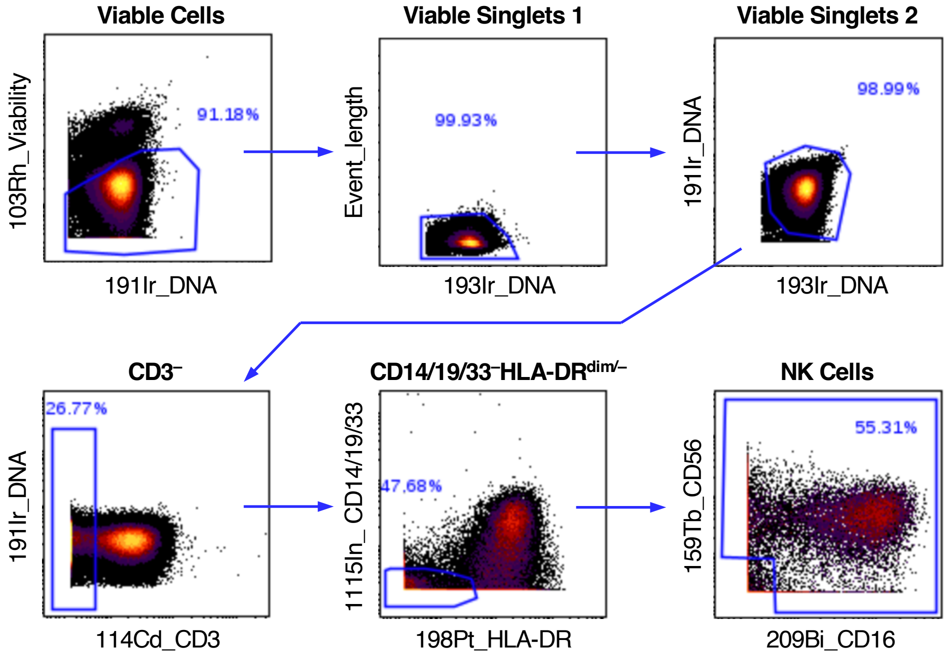


**Supplementary Figure 1. Gating strategy for NK cells on CyTOF sample.** Cryopreserved PBMC were recovered overnight in 10 ng/mL rhIL-15 and profiled by CyTOF following stimulation. Representative FACS plots show manual gating of NK cells prior to unsupervised clustering and downstream analyses.

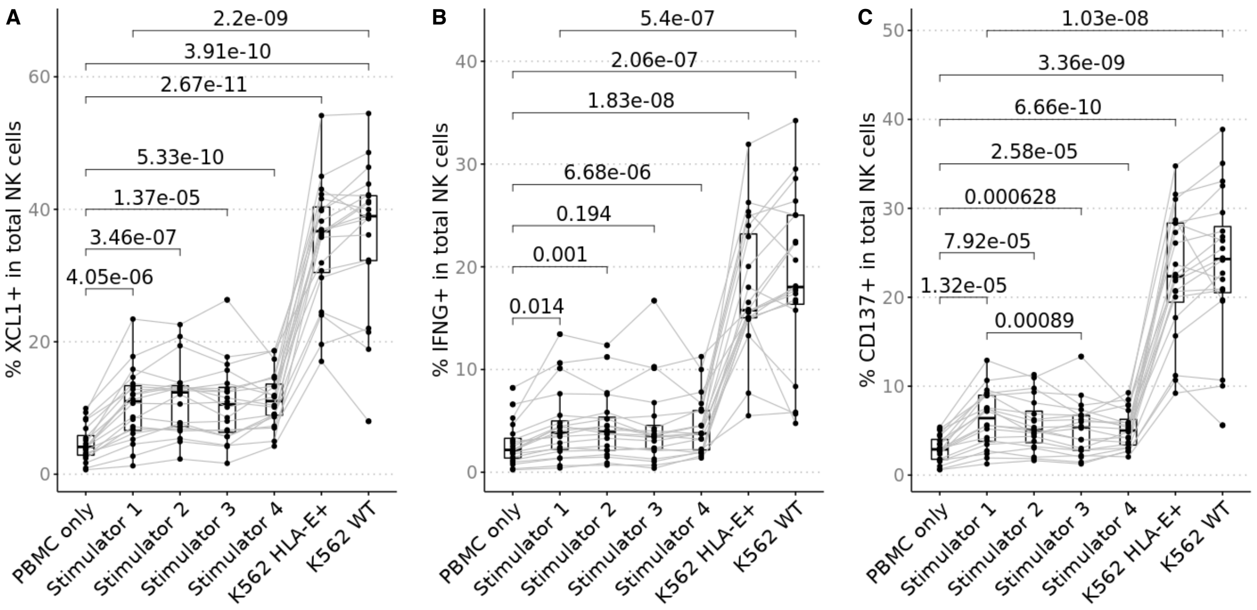


**Supplementary Figure 2. Activation of alloreactive NK cells varies between healthy donors and stimulator cell.** Cryopreserved PBMC from healthy donor (n=20) were recovered overnight in 10 ng/mL rhIL-15 and stimulated for 6 hours 3:1 E/T with 4 distinct allo-stimulator lines, K562 WT and K562 HLA-E+. Results were profiled by CyTOF. Percentage of (A) XCL1+, (B) IFNγ+, (C) CD137+ NK cells across healthy donors increases in response to stimulators. There were no differences between Stimulators 1-4 except for %CD137+ between Stimulator 1 and Stimulator 3. All Stimulators induced less activation than K562 WT and K562 HLA-E+, and there was no difference between K562 WT and K562 HLA-E+. *P* value was calculated using 2-sided paired Student's *t*-test and values were adjusted for multiple testing with Bonferroni correction.

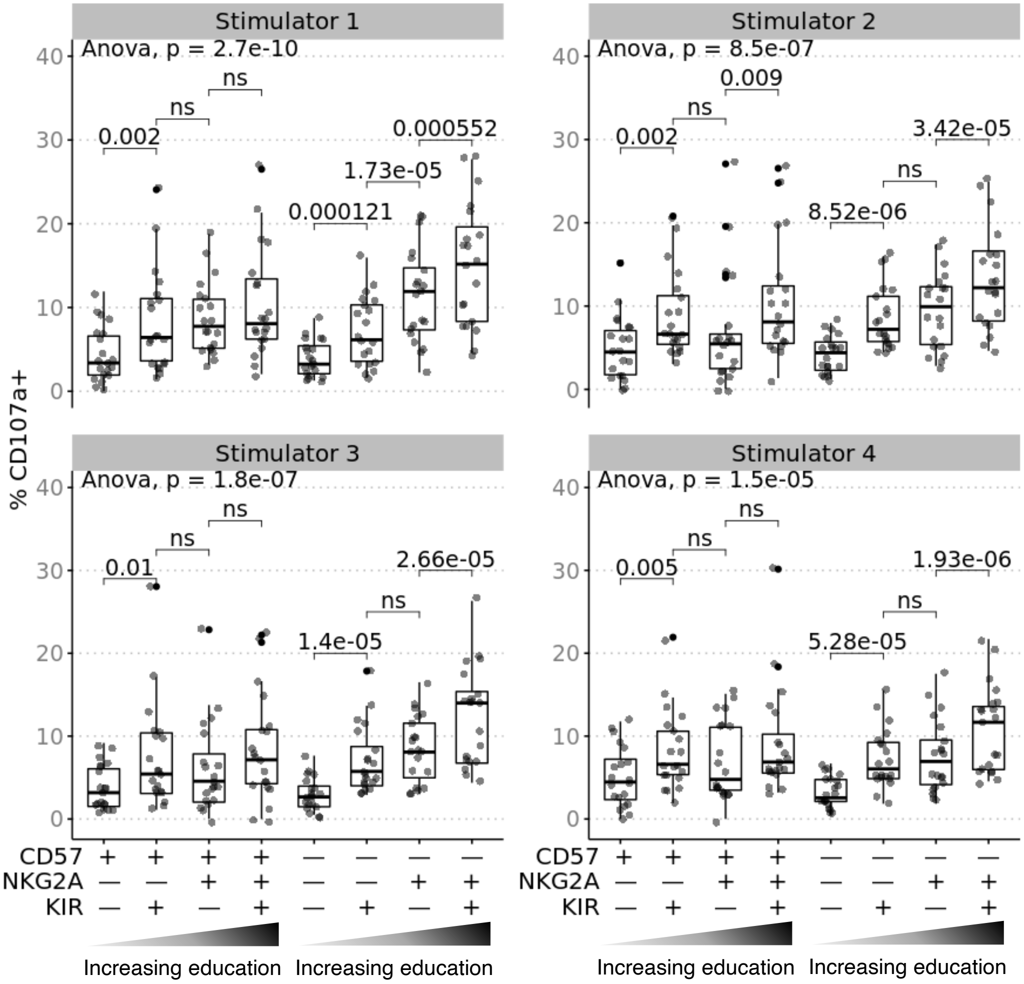


**Supplementary Figure 3. Educated NK cell subsets are more responsive to missing-self when stimulated with all allo-stimulator cell lines.** Cyropreserved PBMC from healthy donor (n=20) were recovered overnight in 10 ng/mL rhIL-15 and stimulated for 6 hours 3:1 E/T with 4 distinct allo-stimulator lines, K562 WT and K562 HLA-E+. Results were profiled by CyTOF. NK cell subsets were defined by gating on combinatorial expression of educating inhibitory receptors, NKG2A, KIR3DL1, KIR3DL2, KIR2DL1/S1, KIR2DL2 and KIR2DL3 and the CD57 maturation marker. NKG2A-expressing and KIR-expressing subsets produce more CD107a in response to B cell allo-stimulators 1-4. *P* value was calculated using 2-sided paired Student's *t*-test and values were adjusted for multiple testing with Bonferroni correction; ns indicates $p > 0.1$.

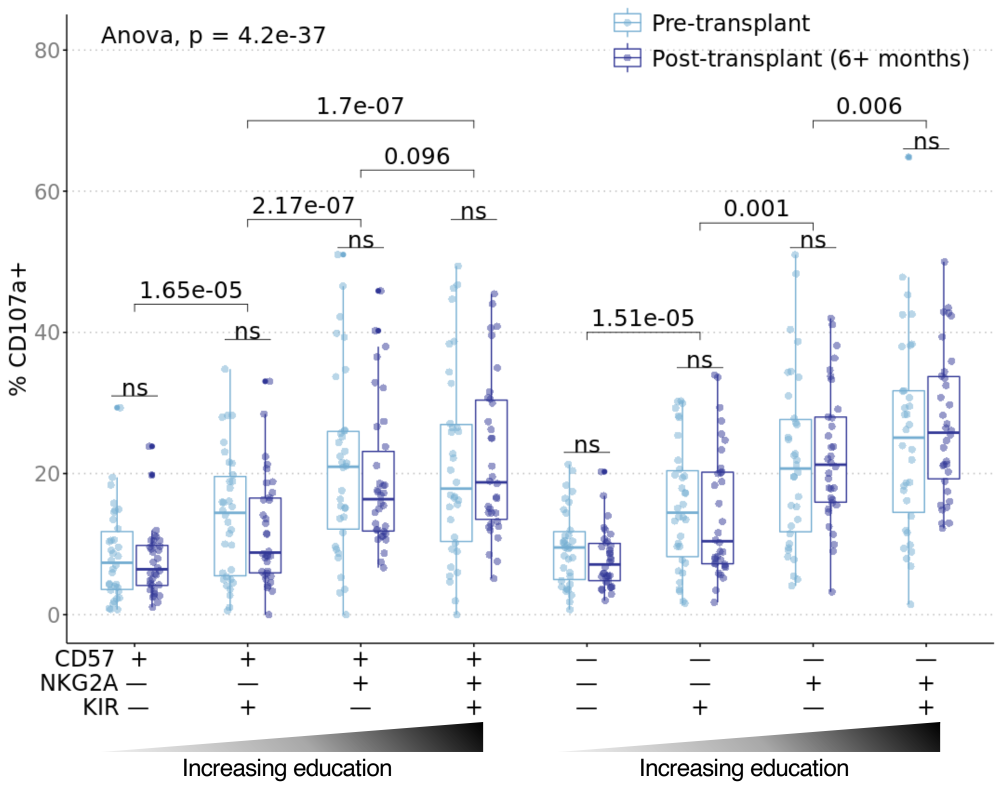


**Supplementary Figure 4. Alloreactivity of educated NK cells transcend donor differences and is maintained post-transplant.** Cryopreserved pre- and post-transplant PBMC from CTOT01 kidney transplant (n=70) were recovered overnight in 10 ng/mL rhIL-15 and stimulated with donor allo-stimulator B cells for 6 hours at 3:1 E/T. Results were profiled by CyTOF and subsets were defined by gating on $CD56^{dim}$ NK cells followed by combinatorial expression of educating inhibitory receptors, NKG2A, KIR3DL1, KIR3DL2, KIR2DL1 and KIR2DL3 and the CD57 maturation marker. Uneducated NK cells, defined as $KIR^{+}$ NK cells in recipients without HLA that encode cognate ligand, were removed from the total $KIR^{+}$ population of that recipient. The increase in alloreactivity of the educated subsets compared to uneducated subsets in the pre- and post-transplant timepoints are consistent with the results in Figure 5. *P* value was calculated as one-way ANOVA as indicated and as 2-sided paired Student's *t*-test; *t*-test *p* values were adjusted for multiple testing with Bonferroni correction and ns indicates $p > 0.1$.

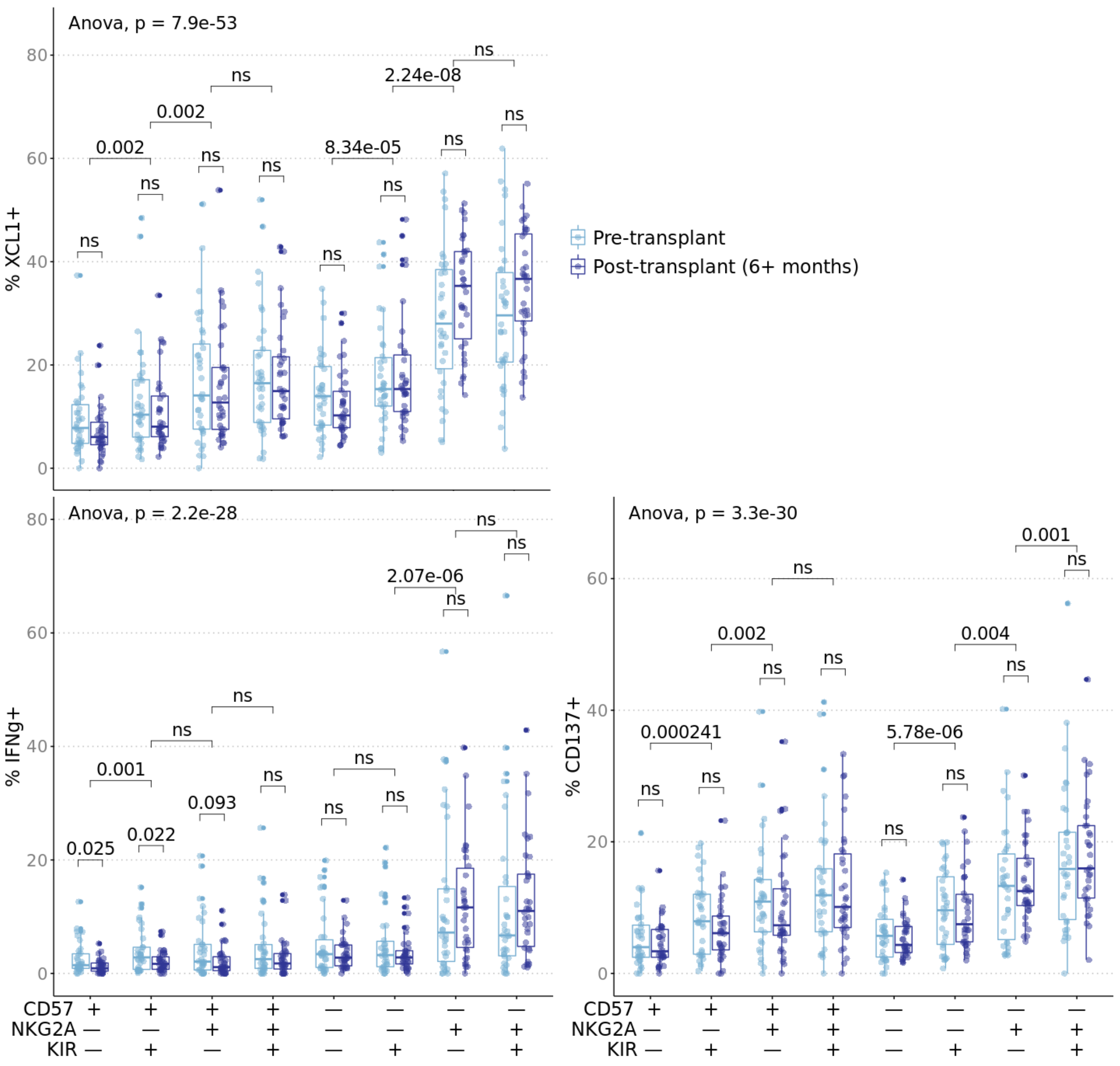


**Supplementary Figure 5. Educated NK cells maintain alloreactivity post-transplant.** Pre- and post-transplant PBMC from CTOT01 kidney transplant (n=70) were recovered overnight in 10 ng/mL rhIL-15 and stimulated with donor allo-stimulator B cells for 6 hours at 3:1 E/T. Results were profiled by CyTOF and subsets were defined by gating on $CD56^{dim}$ NK cells followed by combinatorial expression of educating inhibitory receptors, NKG2A, KIR3DL1, KIR3DL2, KIR2DL1 and KIR2DL3 and the CD57 maturation marker. Upon stimulation with donor cells, recipient $NKG2A^{+}KIR^{+}$ subsets produced the most XCL1, IFNγ, and CD137 while the uneducated $NKG2A^{-}KIR^{-}$ subsets produces the least XCL1, IFNγ, and CD137. The effect of NKG2A/KIR education on CD137 and CD107a expression in response to donor cells persisted 6+ months post-transplant (n=34). IFNγ production in $CD57^{+}NKG2A^{-}KIR^{-}$ and $CD57^{+}NKG2A^{-}KIR^{+}$ NK cells decreased post-transplant. *P* value was calculated as one-way ANOVA as indicated and as 2-sided paired Student's *t*-test. *t*-test *p* values were adjusted for multiple testing with Bonferroni correction and ns indicates $p > 0.1$.

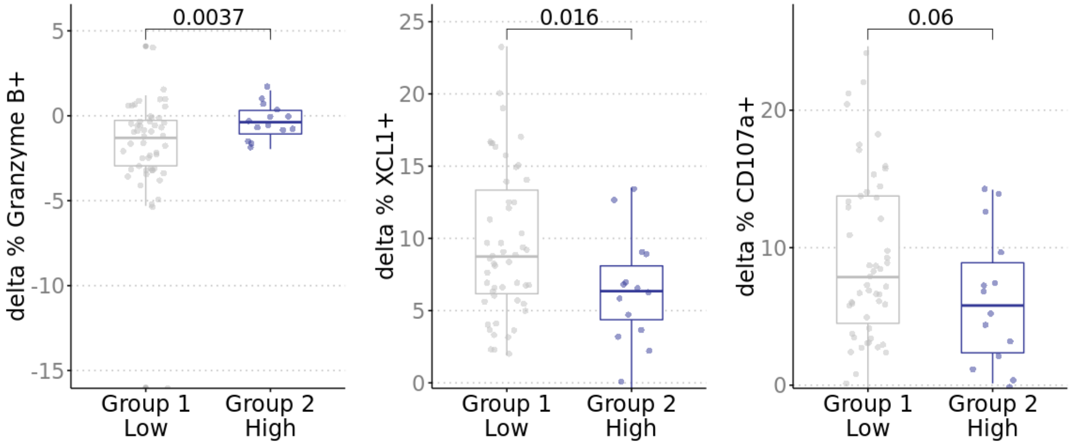


**Supplementary Figure 6. High HLA-E expression on donor cells inhibit recipient NKG2A⁺NKG2C⁻ NK cells.** Donor stimulator groups were defined by expression of HLA-E where 75th percentile was high (n=16) and remaining were low (n=51). Hierarchical clustering defined 14 donors with high HLA-E expression and similar expression of other inhibitory/activating NK ligands and 2 donors with high HLA-E expression that also expressed higher CD112/HLA-F expression compared to other high HLA-E donors. One donor expressed CD112 at 3.3 standard deviations greater than the mean and the other donor expressed HLA-F at 1.8 standard deviations from mean. These two donors were excluded from this analysis. Change in percent positive of CD107a, XCL1 and Granzyme B in NKG2A⁺NKG2C⁻ cells from baseline was greater when recipient cells were stimulated by donors with lower HLA-E. Nominal *p* value was calculated using 2-sided unpaired Student's *t*-test.

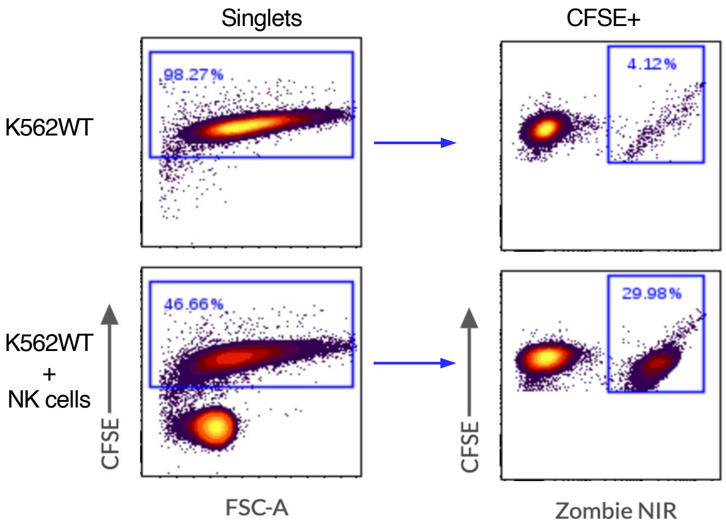


**Supplementary Figure 7. Gating strategy for NK cell killing assays.** PBMC from healthy donor buffy coats were isolated by Ficoll density centrifugation and rested overnight in 10 ng/mL IL-15. PBMCs were enriched for NK cells by negative selection and flow-sorted for $NKG2A^{+}KIR^{+}NKG2C^{-}$, $NKG2A^{+}KIR^{-}NKG2C^{-}$, $NKG2A^{-}KIR^{+}NKG2C^{-}$, and $NKG2A^{-}KIR^{-}NKG2C^{-}$ subsets. NK cell subsets were stimulated with CFSE-stained K562 wildtype and allo-stimulator B cells. Representative FACS plots show gating on $CFSE^{+}$ stimulator cells. Zombie NIR viability dye staining identifies stimulator cells that have died in co-culture with NK cells.
